# Germline DNA replication shapes the recombination landscape in mammals

**DOI:** 10.1101/2020.09.23.308874

**Authors:** Florencia Pratto, Kevin Brick, Gang Cheng, Gabriel Lam, Jeffrey M. Cloutier, Daisy Dahiya, Stephen R. Wellard, Philip W. Jordan, R. Daniel Camerini-Otero

## Abstract

Genetic recombination generates novel trait combinations and understanding how recombination is distributed across the genome is key to modern genetics. The PRDM9 protein defines recombination hotspots, however megabase-scale recombination patterning is independent of PRDM9. The single round of DNA replication, which precedes recombination in meiosis, may establish these patterns, therefore we devised a novel approach to study meiotic replication that includes robust and sensitive mapping of replication origins. We find that meiotic DNA replication is distinct; reduced origin firing slows replication in meiosis and a distinctive replication pattern in human males underlies the sub-telomeric increase in recombination. We detected a robust correlation between replication and both contemporary and ancestral recombination and found that replication origin density coupled with chromosome size determines the recombination potential of individual chromosomes. Our findings and methods have far-reaching implications for understanding the mechanisms underlying DNA replication, genetic recombination, and the landscape of mammalian germline variation.

## Introduction

Sexual reproduction uses a specialized cell division called meiosis, in which a single round of DNA replication is followed by two cell divisions to create haploid gametes. Genetic recombination in meiosis assures faithful segregation of chromosomes and establishes patterns of genetic linkage and inheritance. Recombination is initiated by the formation of hundreds of programmed DNA double-strand breaks (DSBs). In mice and humans, DSBs are targeted by DNA sequence-specific binding of a meiosis-specific histone methyltransferase, PRDM9 (Baudat et al., 2010; Myers et al., 2010; Parvanov et al., 2010). Hundreds of PRDM9 alleles exist (Berg et al., 2010; Buard et al., 2014) that primarily differ in the DNA-binding domain. Thus, each variant may yield a distinct patterning of meiotic DSBs (Smagulova et al., 2016). Nonetheless, megabase-scale similarities between individuals demonstrate that a broad-scale, PRDM9-independent layer of control also shapes meiotic recombination (Davies et al., 2016; Myers et al., 2005; Smagulova et al., 2011). The clearest manifestation of this is seen in the elevation of meiotic DSBs in subtelomeric DNA of human males, independent of PRDM9 genotype (Pratto et al., 2014).

Since DNA replication determines genome structure (Klein et al., 2019) and immediately precedes DSB formation in meiosis, we hypothesized that DNA replication may drive the broad-scale regulation of recombination in mammals. Replication and recombination are correlated in yeasts (Borde et al., 2000; Murakami and Nurse, 2001) and in barley (Higgins et al., 2012). A mechanistic link has been demonstrated in baker’s yeast, where a component of the DSB machinery is activated by passage of the replication fork (Murakami and Keeney, 2014). This results in a spatio-temporal coordination between replication and DSB formation.

All current knowledge of DNA replication in mammalian meiosis stems from classical papers using early molecular and cytological techniques (for review see (Chandley, 1986)). The paucity of studies of DNA replication in mammalian meiosis stems from the requirement to study meiosis *in-vivo*, in the context of a complex tissue (Handel et al., 2014). This contrasts with cell-types that can be studied *in vitro*; indeed, mammalian DNA replication is exclusively studied in cell culture. This presents difficulties in adapting techniques for studying replication, which have been mostly designed and optimized for cell culture. To address this shortcoming, we devised the first method to map origins of replication in mammalian tissue (in this case, testis), developed a cell-type specific method to interrogate replication timing in meiotic S-phase and designed an *in-silico* modelling strategy to parameterize DNA replication *in-vivo*. This unique tripartite approach has generated the first comprehensive, parameterized description of DNA replication genome-wide in meiosis, or, for that matter, in any mammalian tissue. We found that DNA replication in the germline is distinct from replication in other cell types and that this unique replication patterning plays a direct role in shaping megabase-scale patterns of meiotic recombination and genome diversity.

## Results

### Highly specific replication origin mapping in mammalian testis

To identify origins of replication, we introduced an important adaptation to an existing method to sequence the RNA-primed short nascent leading strands (SNS) (Bielinsky and Gerbi, 1998) (Figure 1A). Briefly, RNA-primed leading strands are isolated by using lambda exonuclease to digest all DNA that lacks an RNA primer. Okazaki fragments, while also RNA-primed, are excluded by size selection. A key improvement upon SNS-Seq (Cayrou et al., 2012; Fu et al., 2014; Jodkowska et al., 2019; Picard et al., 2014) is that we directly sequence the captured single-stranded DNA (Khil et al., 2012) at origins. This retains the strand information of the nascent strand DNA (Origin-derived Single-Stranded DNA Sequencing; Ori-SSDS) (Figure 1A) and avoids biases resulting from random-priming and second-strand synthesis (Sequeira-Mendes et al., 2019). Characteristic bidirectional replication at origins manifests as a strand switch in sequenced reads. Distinct from SNS-Seq experiments, we can differentiate true origins from non-specific signals (Figure 1A-C, S1, S2) by imposing a requirement for leading-strand asymmetry at origins (see methods). Accounting for strand asymmetry substantially reduces the number of replication origins estimated by SNS-Seq experiments (40-50%; Figure S3; (Cayrou et al., 2015)) because local peaks within SNS-Seq signals have been spuriously defined as very closely-spaced replication origins (Figure S3).

**Figure 1.**
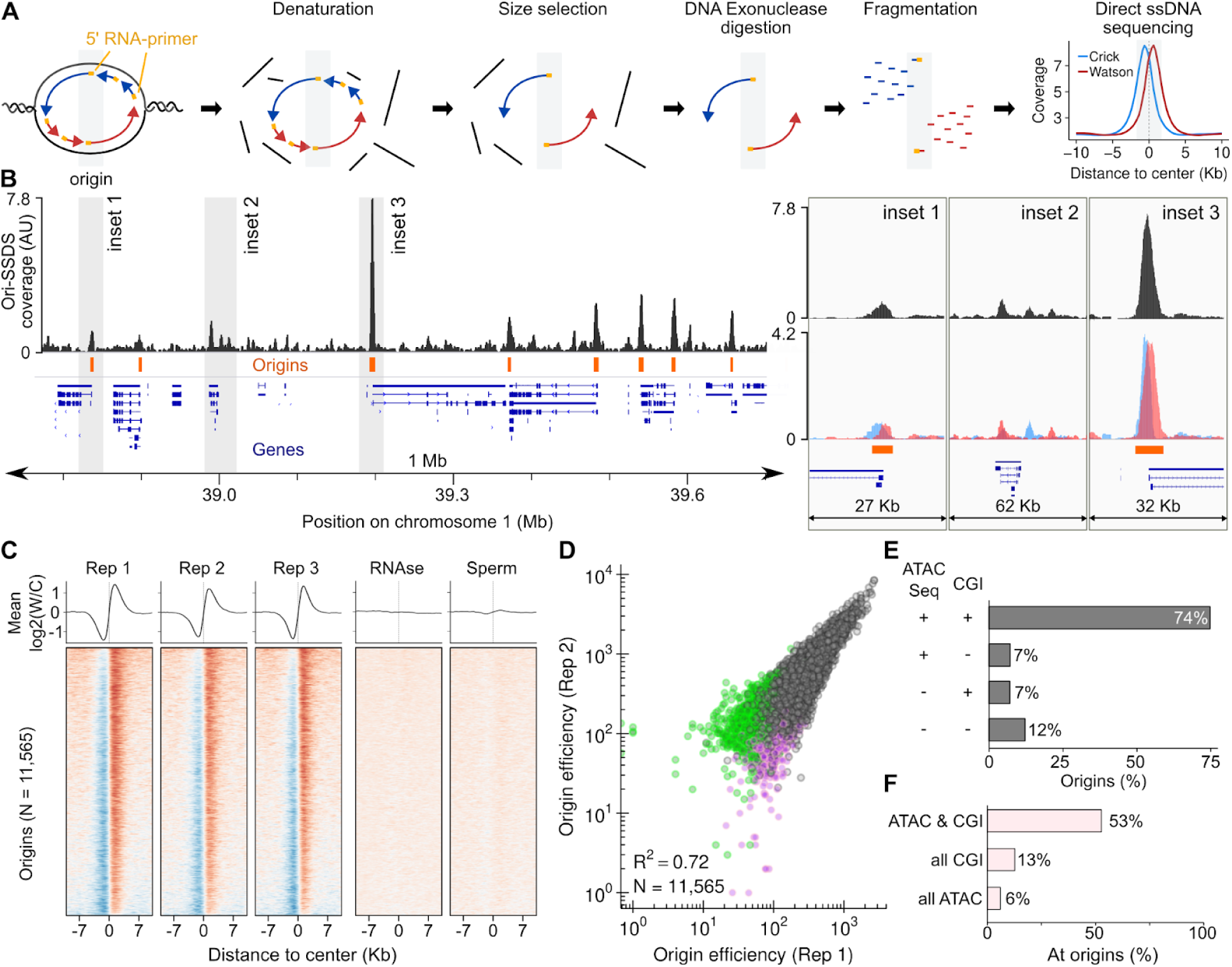
Identification of replication origins. **(A)** Schematic of the Ori-SSDS protocol to sequence nascent leading strands. **(B)** The Ori-SSDS signal in a typical 1 Mb region. Insets show the strand-specific Ori-SSDS signal at representative weak (inset 1) and strong (inset 3) origins and at a series of non-origin peaks (inset 2) that lack Ori-SSDS Watson-Crick asymmetry; red = Watson-strand coverage, blue = Crick-strand coverage). Coverage is calculated in 1 Kb windows with a 147 bp step. **(C)** Ori-SSDS signal at origins is reproducible but is lost in control experiments. The log_2_ ratio of Watson/Crick strand ssDNA fragments is shown, smoothed in 1 Kb windows. **(D)** Origin efficiency is highly correlated among replicates. Origin efficiency varies ∼100-fold. Origins found only in one replicate are colored purple and green, respectively. **(E)** Origins of replication occur predominantly at CpG islands (CGIs) that coincide with open chromatin as measured by ATAC-Seq (Maezawa et al., 2018). **(F)** Origins are only detected at about half of the CGIs at ATAC-Seq peaks.

**Figure 2.**
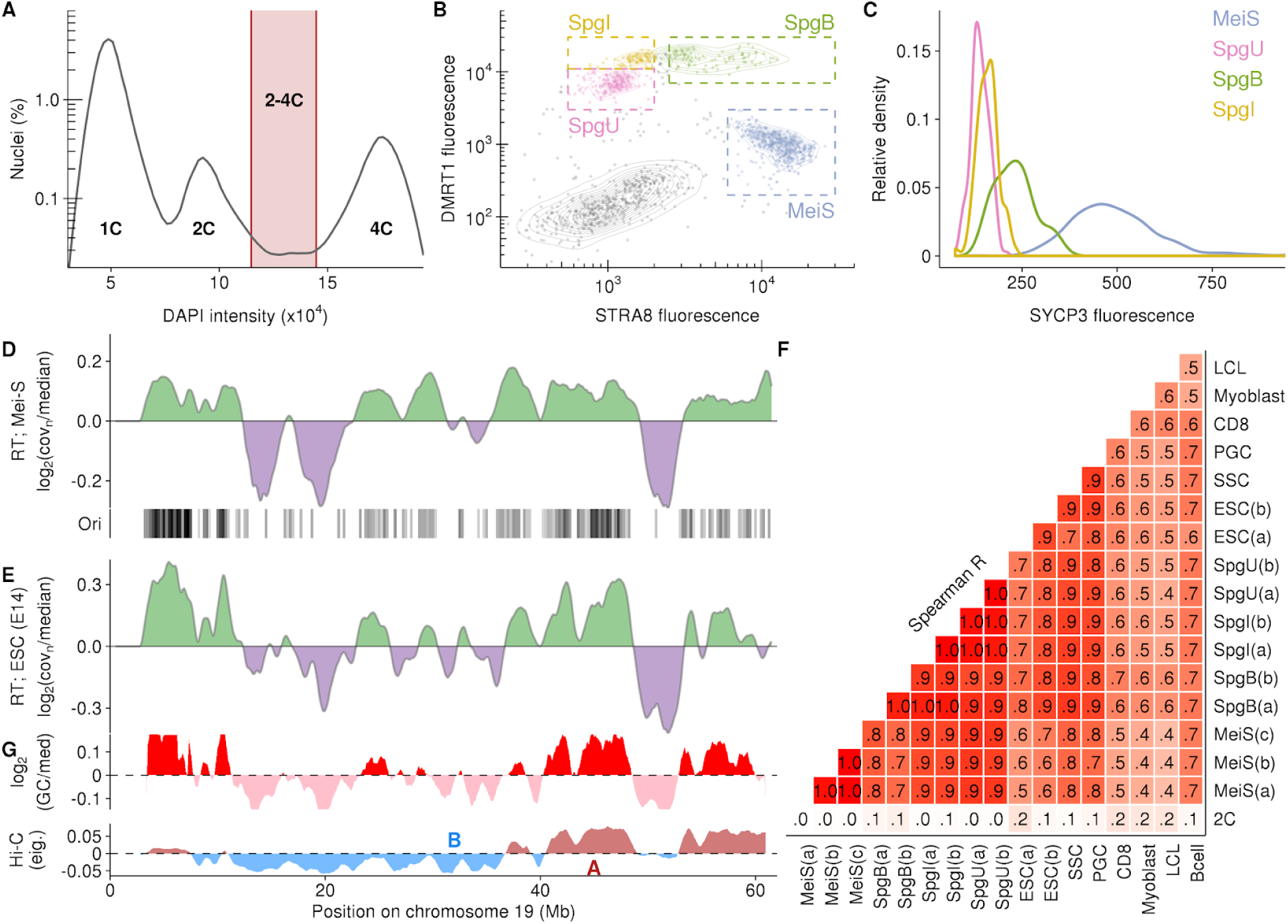
Replication timing in meiotic prophase I of male mice. **(A-C)** Meiotic S-phase and spermatogonia nuclei were isolated using fluorescence-activated nuclei sorting (Lam et al., 2019). **(A)** Replicating nuclei were isolated by gating 2-4C nuclei by DAPI content. **(B)** Meiotic S-phase nuclei (MeiS) can be distinguished from S-phase nuclei of pre-meiotic germ cells using positive selection for nuclei expressing STRA8, and negative selection for DMRT1. This sorting paradigm can also distinguish three sub-types of germ cells (undifferentiated spermatogonia, SpgU; intermediate spermatogonia, SpgI; type-B spermatogonia, SpgB) **(C)** SYCP3 is a meiosis-specific protein that was not used for nuclei sorting. SYCP3 is elevated in the isolated meiotic nuclei relative to the other populations. Above-background SYCP3 is also seen in the SpgB population. These are the germ-cells that immediately precede meiotic entry. **(D)** Replication timing (RT; log_2_ of normalized sequencing coverage (cov_n_)/genome median) in meiotic S-phase nuclei correlates with origin locations in testis. **(E)** RT in E14 embryonic stem cells. **(F)** RT is highly correlated across cell-types. RT was inferred from published whole genome sequencing data for Spermatogonial Stem Cells (SSC), Primordial Germ Cells (PGC), Embryonic Stem Cells (ESC) CD8^+^ cells (CD8), and activated B-cells (Bcell). Pre-processed RT data was obtained for Myoblast and Lymphoblastoid Cell Lines (LCL). Details of samples in Tables S1-3. (**G)** GC-content (in 10Kb windows; shown as log_2_(GC/meanGC); smoothed in 1Mb windows, 10Kb steps) and genome compartmentalization at zygonema (Patel et al., 2019) also correlate with RT. Hi-C track shows the eigenvector values for the first principal component of the Hi-C matrix (calculated in 100 kb windows, smoothed in 1Mb windows, 10Kb steps). Active (A) and inactive (B) compartments are highlighted.

We detected 10,932, 12,100 and 13,082 origins in replicate experiments from testes of individual male mice, demonstrating that our method is sufficiently sensitive to derive replication origin maps from individual animals. Prior to our experiments, the number of cells required to detect origins appeared to be prohibitive for *in-vivo* studies; for example, SNS-Seq typically starts from 10^9^ exponentially growing cultured cells (Almeida et al., 2018). We used just 2 x 10^6^ replicating cells (Kojima et al., 2019) for Ori-SSDS; this is 3-5 fold fewer than methods that immunoprecipitate pre-replication complexes (i.e. ORC1/ORC2 ChIP-Seq (Miotto et al., 2016)) and orders of magnitude fewer than are required for the sequencing of Okazaki fragments (Petryk et al., 2016). 94% of origins in the smallest set were found in at least one of the other two experiments (Figure S4) and origins unique to individual samples were weak (Figure 1D, S5). No origins were detected in control experiments, either using non-replicating tissue (sperm) or by hydrolyzing the leading strand RNA-primer (see above and methods) (Figure 1C, S1). 94% of replication origins detected by Ori-SSDS in testis are found in SNS-based origin maps from other cell types (Mouse Embryonic Fibroblasts (MEFs) and Embryonic Stem (ES) cells; Figure S6). Furthermore, data from Okazaki-fragment sequencing (in activated B-cells) shows a switch in the polarity of Okazaki fragments around origin centers (Figure S6). These data strongly suggest that Ori-SSDS accurately captures origins of replication from tissue and that origins in testis mostly occur at sites that are also used as origins in other cell types.

**Figure 5.**
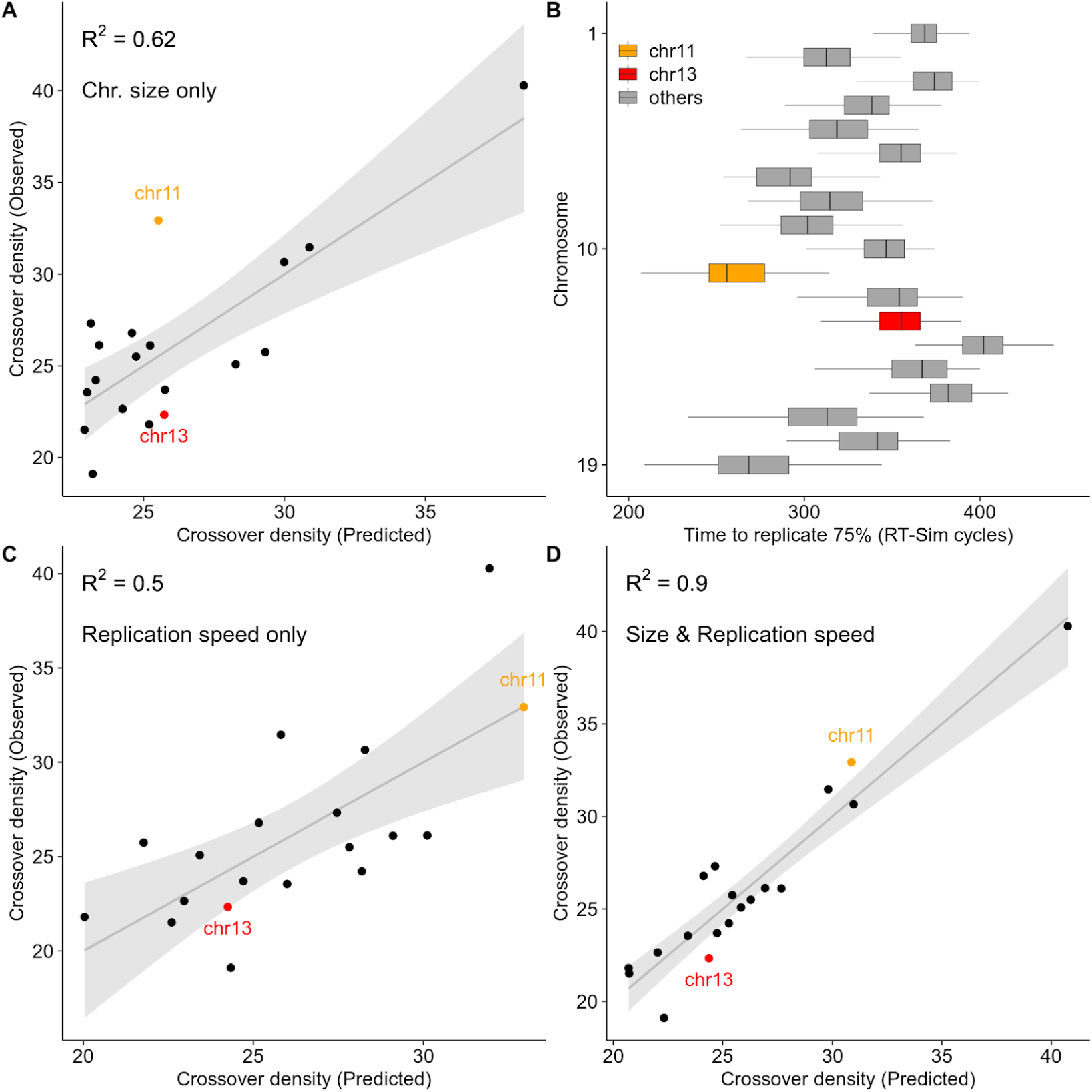
DNA replication predicts per-chromosome crossover rates. **(A)** A linear regression model can predict crossover density from chromosome size (polynomial fit). Crossover density is calculated as total crossovers per Mb. Crossovers from (Yin et al., 2019). R^2^ is strongly influenced by the single small outlier chromosome (chr19; rightmost). **(B)** Chromosomes replicate asynchronously. Chromosome 11 and 13 are highlighted because while they are a similar size, chromosome 11 replicates far earlier. The time required to replicate 75% of each chromosome is inferred from the best-fitting RT-Sim model (see also Figure S16). Boxplots indicate the range of times from individual modelled cells. **(C)** A linear regression model can predict crossover density from replication speed. **(D)** Per chromosome crossover density can be accurately predicted from a linear model that combines replication speed and chromosome size.

Origin efficiency is a measure of the frequency with which an origin is used. We use the sequencing depth of “correctly” oriented Ori-SSDS read-pairs to infer origin efficiency (see methods). This efficiency varies ∼100-fold and is highly correlated between replicates (Figure 1D, S3; Spearman R^2^ = 0.65-0.74). Despite the fragment asymmetry around the origins, there is considerable overlap of the Watson and Crick signals (Figure 1A, S1). Thus, while DNA replication originates at discrete loci, the initiation window spans several hundred base-pairs at the origin center. This contrasts with *S. cerevisiae*, where an 11 bp motif defines a more precise origin of replication (Broach et al., 1983; Stinchcomb et al., 1980). Together, these data show that Ori-SSDS can identify the origins of replication genome-wide from mammalian tissue. 11,565 high-confidence origins, found in at least two of three replicates (Figure S4, S5), were used for subsequent analyses.

**Figure 3.**
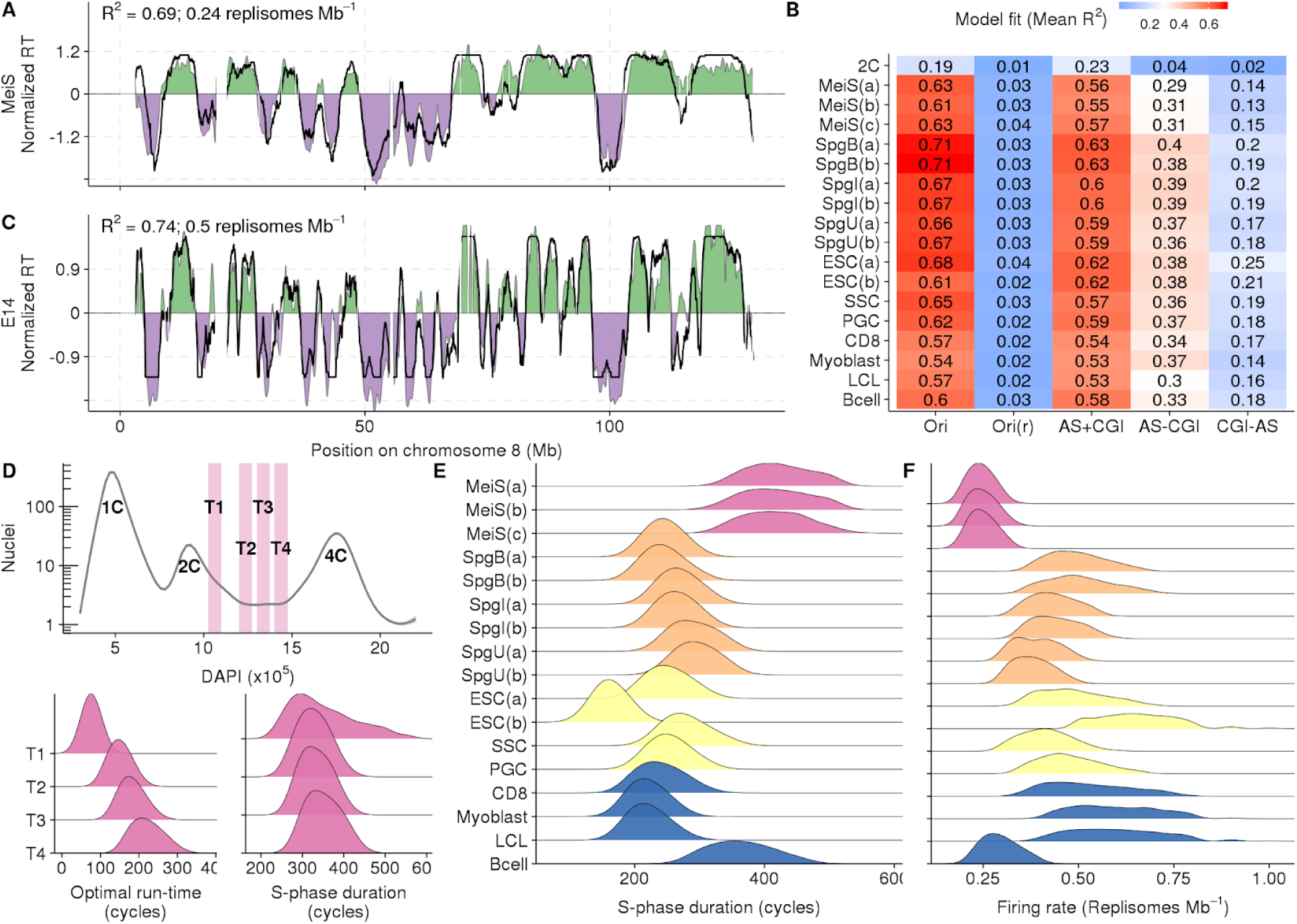
*In silico* modeling recapitulates RT from origin locations. **(A)** Best fitting RT-Sim model for Meiotic S-phase RT (MeiS(a)). Experimentally determined RT is shown as filled area. Simulated RT is the black line. Both simulated and experimental RT are normalized by mean and standard deviation (s.d.) (RTnorm = (RT-mean(RT)) / s.d.(RT)). **(B)** Fit scores for models built using RT from different cell types and different datasets as a proxy for origins. Mean of the top 0.015% of models is shown. Samples are described in detail in Tables S1-3. Briefly, RT is from: MeiS = Meiotic S-phase, Spg(B,I,U) = B-type, Intermediate and Undifferentiated Spermatogonia, ESC = Embryonic Stem Cells, SSC = Spermatogonial Stem Cells, PGC = Primordial Germ Cells, CD8 = CD8 cells, Myoblast = Myoblast cell line, LCL = Lymphoblastoid cell line, Bcell = Activated B-cells. (a,b,c) designate replicates. ESC(a) is from published whole genome sequencing data, ESC(b) is from cells grown by us. Ori = OriSSDS origins, Ori(r) = OriSSDS origins with randomized genomic location, AS+CGI = ATAC-Seq peak at a CGI, AS-CGI = ATAC-Seq peak not at a CGI, CGI-AS = CGI not at an ATAC-Seq peak. **(C)** Best fitting RT-Sim model for RT in E14 ESCs (ESC(a)). **(D)** Sorting schematic for S-phase populations in lower panel. Populations have increasing DNA content from T1 to T4. The optimal simulation run-time for best-fitting models correlates with increasing DNA content. Nonetheless, the predicted S-phase duration is similar among all populations. **(E)** Total replication time for best-fitting models. Meiotic S-phase is notably slow. **(F)** Best fitting models imply that meiotic S-phase is slow because fewer replisomes fire than in other cell types. Activated B-cells appear to share this property of replication with meiotic S-phase.

Origins identified from mouse testis are unevenly distributed in the genome (Figure S7A) and cluster in gene rich regions; 66% of origins occur within 1 Kb of a transcription start site (TSS) (Figure S7B). DNA at the origin center is intrinsically flexible, flanked by relatively rigid DNA, and this may reflect a requirement of the ORC complex to bend DNA (Li et al., 2018) (Figure S7C). Origins are GC-rich, with elevated CpG, and to a lesser extent, GpC dinucleotides (Figure S7D). GC content, CpG density and GpC density are all positively correlated with origin efficiency (Figure S7E) implying that nucleotide content either directly or indirectly plays a role in origin firing. G-quadruplexes, a predicted secondary structure implicated in origin firing (Valton et al., 2014), are also strongly enriched at origin centers (Figure S7F). 74% of origins coincide with CpG islands (CGIs) in accessible chromatin (as measured by ATAC-Seq in spermatogonia - the germ-cells that precede meiosis (Maezawa et al., 2018); Figure 1E), yet only 53% of such sites are used for origin formation (Figure 1F). CGIs in inaccessible chromatin and open chromatin lacking a CGI are far less predictive of origin locations (Figure 1E,F). We did not identify any conserved sequence motif at replication origins, however the density of CpG dinucleotides alone can predict origin locations (Figure S7G). Indeed, CpG density is a better predictor of origin locations than binding sites are for transcription factors (Figure S7H). Many of these observations are consistent with properties of replication origins defined in cultured cells (Marchal et al., 2019; Miotto et al., 2016), and together suggest that replication in the germline initiates preferentially at CpG islands, near gene promoters, in accessible chromatin.

Broad replication initiation zones - up to 150 Kb and comprising ∼7% of the genome - have been proposed as a major mode of replication initiation in humans (Petryk et al., 2016). Although most origins mapped by Ori-SSDS are narrow and discrete, 1,925 Ori-SSDS peak are wider than 10Kb, and may represent replication initiation zones (the expected width of the Ori-SSDS signal is approximately ± 3 Kb around the origin of replication; determined by the size selection step; see methods; Figure 1A; Figure S8A) To assess if putative initiation zones result from closely-spaced, but difficult-to-resolve discrete origins, we examined the Ori-SSDS signal around local peaks in CpG density (CpG peaks coincide with the center of isolated origins; Figure S8B). We found that within our defined initiation zones, most local CpG peaks exhibited origin-like Ori-SSDS strand asymmetry, implying that these sites are *de-facto* origins (Figure S8C). 620 zones contained multiple CpG peaks (N_CpG Peaks_ = 1,645), thus, closely-spaced and difficult-to-resolve origins account for a large number of initiation zones (Figure S8D-F). Among such zones is the extensively studied *HoxA* locus, where Ori-SSDS data suggests that ∼19 origins of replication occur within just 110 Kb (Figure S8E). 816 zones contained a single CpG peak, and likely represent origins where we captured longer-than-expected nascent strands (Figure S8G). The remaining 490 zones did not contain any CpG peak above our threshold and may represent more amorphous replication initiation regions. Alternatively, they may represent clusters of weak origins (Figure S8G,H). Overall, we found extensive evidence that clustered, yet discrete origins of replication can be mistaken for initiation “zones”. Previous experiments that identified “zones” lacked the resolution to detect this fine sub-structure.

### An unambiguous snapshot of replication in meiotic S-phase

Stochastic origin firing and uneven origin density result in distinct earlier and later replicating parts of the genome. We inferred this replication timing (RT) in meiosis. Meiotic S-phase nuclei constitute just 1% of cells in adult testis (Kojima et al., 2019). Using a variant of fluorescence activated nuclei sorting (Lam et al., 2019), we isolated up to 2 x 10^5^ meiotic S-phase nuclei at 94% purity (inspection of wide-field images; Figure S9) from a single adult mouse (Figure 2A-C); this relied on a combination of DNA content, negative selection for a marker of non-meiotic cells (DMRT1) and positive selection for a meiotic protein (STRA8). A marker not used for sorting, but with expected meiotic expression (SYCP3) validated the purity of the meiotic population (Figure 2C). We then inferred meiotic RT from coverage imbalances in whole-genome sequencing of these nuclei (see methods; RT-Seq) (Koren et al., 2014). Replication origin density is high in early replicating regions and low in late-replicating DNA (Figure 2D). This yields the first snapshot of replication timing in mammalian meiosis. We also inferred RT from the other major replicating cell types in adult testis - undifferentiated and differentiating spermatogonia - and from published whole genome sequencing (WGS of other cell types (see methods; Figure S10). Meiotic RT profiles are highly correlated among replicates and, to a lesser extent, with RT from other cell types (Figure 2D-F, Figure S10). In particular, meiotic RT is notably similar to RT in the cells that immediately precede meiotic S-phase and to RT in other germ cells (PGCs, SSCs, ESCs). RT in all cell-types is highly correlated with GC-content (Spearman R^2^ = 0.27 - 0.40) and with nuclear compartments (Spearman R^2^ = 0.57 - 0.70)(Figure 2G).

### *in-silico* modeling recapitulates replication timing from origin locations

RT-Seq yields a static snapshot of replication and these static RT-Seq snapshots from different cell-types are often remarkably similar. Furthermore, the differences between RT-Seq profiles are difficult to interpret because RT-Seq can be strongly influenced by the average stage of S-phase progression and by the relative synchrony of the population (Figure S11). We hypothesized that subtle differences in the properties of replication among cell types could be captured by *in-silico* modelling.

We built an *in silico* model of DNA replication that required experimentally defined origins as input and that outputs a simulated RT profile. By examining the properties of replication that produce a best-fit between simulated RT and experimental RT, we can understand the parameters of replication that yield different RT profiles. Modeling parameters are the number of active forks, whether to use origin efficiency estimates as a firing probability, and the total duration of simulated S-phase (see methods). Replication fork speed is assumed constant among cell types and throughout S-phase. After an initial round of simultaneous origin firing, further origins fire only when two extant forks collide; this simulates the presence of factors that limit the number of active origins (replisome ceiling), deemed important by previous simulations of DNA replication (Gindin et al., 2014; Kelly and Callegari, 2019). Best fitting models were obtained using a grid search for optimal model parameters (see methods).

We found that *in silico* modeling could accurately recapitulate experimental RT from both meiotic (Figure 3A,B, Figure S12A) and non-meiotic S-phase cells (Figure 3B,C, Figure S12B). In contrast, we could not obtain good-fitting models for non-replicating cells (2C) or when origin locations were randomized (Ori(r); Figure 3B). Our success in modeling RT in all cell types using testis-derived origins implies that a common set of origins are used during meiotic and mitotic replication. Indeed in yeast, origins of replication are common to meiotic and mitotic cells (Blitzblau et al., 2012; Wu and Nurse, 2014). Most testis-derived replication origins coincide with CpG islands in open chromatin (ATAC-CGIs; Figure 1E,F), and these regions are broadly similar across diverged cell types in mouse (Figure S13). RT can be modeled using ATAC-CGIs as a proxy for origins, but these models are slightly worse than those using *de facto* origins (AS+CGI; Figure 3B). Finally, RT-Sim does not explicitly distinguish between “early” and “late” firing replication origins, suggesting that the paradigm of early and late origins is not required to explain RT. Instead, “early replicating regions” and “late replicating regions” are defined by high and low origin density, respectively. This is consistent with recent studies in other mammalian cell types (Dileep and Gilbert, 2018; Gindin et al., 2014; Miotto et al., 2016; Takahashi et al., 2019).

### Reduced origin firing slows DNA replication in meiosis

To validate that RT-Sim can explain meaningful properties of replication, we examined model run-time of best-fitting models for very early-, early-, middle- and late- S-phase nuclei (Figure 3D). Optimal model runtime - defined as the time at which simulated RT best fits experimental RT - should reflect the time a population has spent in S-phase. Indeed, optimal runtime got progressively longer from very early through late S-phase populations (Figure 3D), demonstrating that modeling yields meaningful numeric insights into the properties of DNA replication. Importantly, the differences between experimental RT in these populations do not substantially affect model estimates of total S-phase duration (Figure 3D). Accurate estimates of the duration of meiotic S-phase in mammals are notoriously difficult to obtain (Kofman-Alfaro and Chandley, 1970); in mice, estimates range from 14 hrs (Monesi, 1962) to 29 hrs (Ghosal and Mukherjee, 1971). Like in other organisms (Cha et al., 2000; Kofman-Alfaro and Chandley, 1970; Lee and Amon, 2001), DNA replication in meiotic S-phase is thought to be longer than S-phase in non-meiotic cells (Kofman-Alfaro and Chandley, 1970). We find that for best-fitting models, median S-phase duration in meiosis is 1.4 - 1.8 fold longer than in spermatogonia - the cells that precede meiosis (Figure 3E, S14), despite having highly correlated experimentally-measured RT (Figure 2F). In *Saccharomyces cerevisiae*, the slow-down of DNA replication in meiosis may be partly to facilitate recombination, as knocking out *Spo11* - the protein that makes meiotic DSBs - reduces S-phase duration by 30% (Cha et al., 2000). Unlike in yeast, we found no reduction in meiotic S-phase duration in *Spo11^-/-^* mice (Figure S14).

The slow-down we observe in meiotic S-phase results from the use of fewer replisomes; best-fitting models use just 0.24 - 0.28 replisomes Mb^-1^ in meiosis (648 - 756 replisomes per haploid genome; 2,700 Mb genome), compared to 0.44 - 0.54 replisomes Mb^-1^ in ES cells (1,200 - 1,500 replisomes per haploid genome) (Figure 3F). These estimates in ES cells are similar to the replisome count in cultured mouse C2C12 cells (imaging-based estimates; 0.46 - 0.48 replisomes Mb^-1^; (Chagin et al., 2016)). The decreased replisome density in meiosis mirrors findings in newt spermatocytes, where replication tracts in meiosis were notably longer than in mitotic cells (Callan, 1973). Although we don’t explicitly model fork speed, universally altering fork speed without changing origin density cannot explain the presence of longer replication tracts in meiosis. Indeed, in other organisms, fork speed does not vary between meiosis and mitosis (Borde et al., 2000; Callan, 1973). By extrapolating from published estimates of S-phase duration in intermediate-stage spermatogonia (12.5 hr (Monesi, 1962)), we estimate that meiotic S-phase in mice takes 21-24 hrs. Interestingly, the range of S-phase duration estimates for the best-fitting models of meiotic S phase is larger than that of the other populations (Figure 3E, S14). Given that the model simulates replication in single cells, this might indicate that there is a higher heterogeneity of RT profiles in meiosis vs mitosis. This is in line with the experimental data spread seen in yeast (Cha et al., 2000).

Together, these analyses suggest both that the origins captured by Ori-SSDS reflect meiotic replication initiation and that the same origins can yield tangibly different RT profiles if other properties of replication vary. This tripartite approach of origin mapping, RT-Seq and *in-silico* modeling, offers an alternative to classical cytogenetic approaches and yields an unprecedented description of the DNA replication landscape.

### Direct coupling of replication with meiotic recombination

Programmed DSB formation in meiosis occurs after DNA replication. In *Saccharomyces cerevisiae*, passage of the replication fork favors meiotic DSB formation, likely through the phosphorylation of Mer2 (a key protein for DSB formation) (Murakami and Keeney, 2014). This interplay is completely unexplored in large mammalian genomes. In mice and humans, local DSB patterning is determined by the sequence-specific binding of PRDM9 (Baudat et al., 2010; Myers et al., 2010), yet at megabase scales, DSB density is highly correlated between individuals with different PRDM9 alleles (Davies et al., 2016; Smagulova et al., 2011). We therefore asked whether DNA replication underlies the megabase-scale control of DSB patterning.

Meiotic DSBs form through a well-documented cascade; PRDM9 tri-methylates H3K4 (Brick et al., 2012; Diagouraga et al., 2018) and/or H3K36 (Diagouraga et al., 2018; Powers et al., 2016) which appear to recruit the DSB complex. A DSB is made by the SPO11 protein, which is then released with a short oligonucleotide (Bergerat et al., 1997; Keeney et al., 1997; Neale et al., 2005). The 5’ DNA is resected and the DMC1 protein loads on exposed ssDNA to facilitate homologous recombination (Bishop et al., 1992). DSBs ultimately repair as a crossover (CO) or as a non-crossover (NCO). H3K4me3 ChIP-Seq, SPO11-associated oligo mapping (Lange et al., 2016), DMC1 ChIP-single-stranded DNA sequencing (Brick et al., 2012) and repair outcome mapping (COs/NCOs) provide differing quantitative readouts of intermediates in this cascade (Figure 4A).

**Figure 4.**
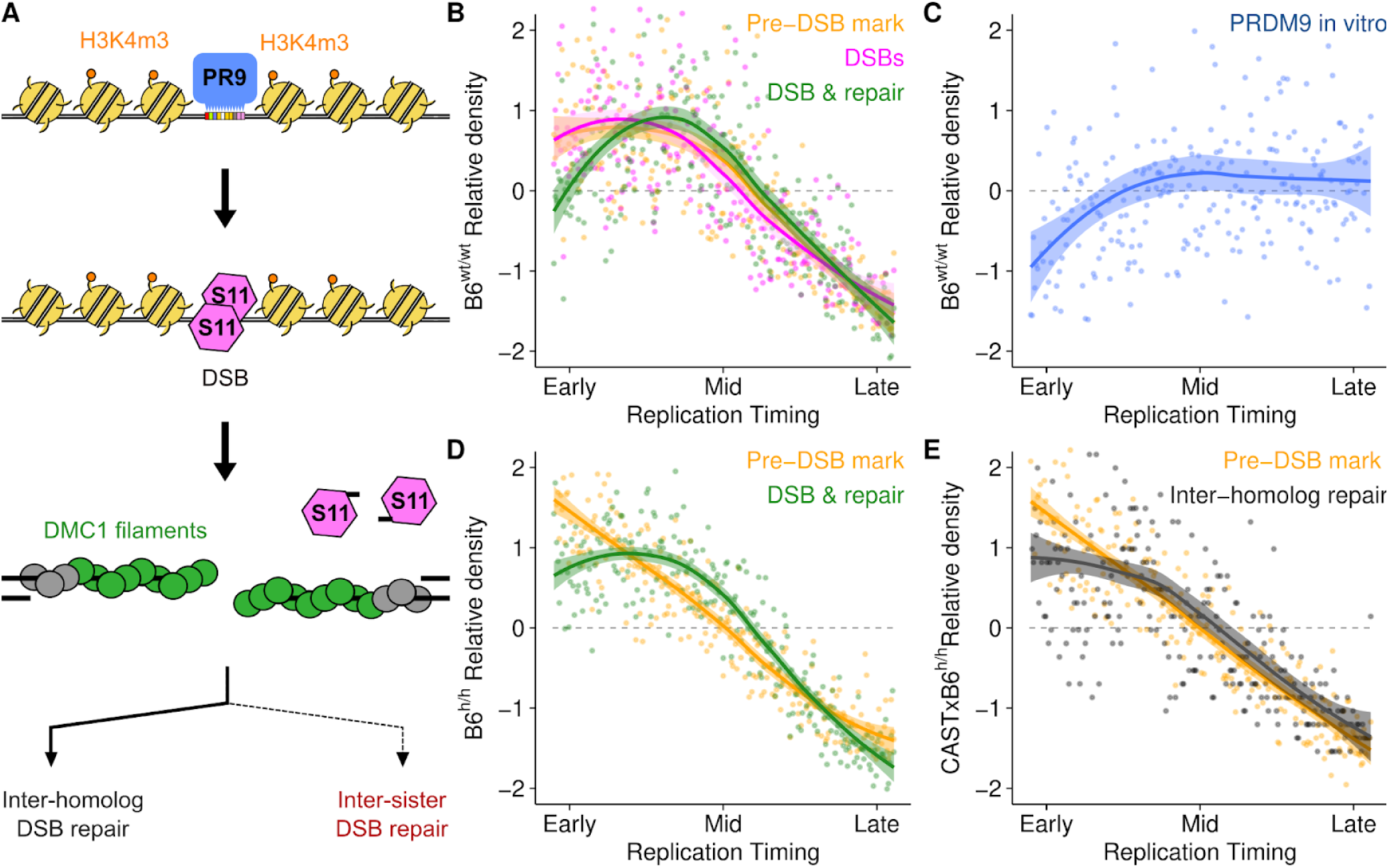
Rapid DSB repair and elevated meiotic recombination in early-replicating regions. **(A)** Schematic of meiotic recombination. The colored lines in B-D reflect the colors in this schematic. PR9 = PRDM9; S11 = SPO11. **(B,D)** PRDM9-mediated H3K4m3 at DSB hotspots (yellow) is enriched in early replicating DNA in both wild-type B6 male mice (B6^wt/wt^; B) and in mice with a humanized *PRDM9* allele (B6^h/h^; C). DSB formation (SPO11-oligo mapping; pink) follows this Pre-DSB mark (B). The DMC1-SSDS (green) signal decays relative to H3K4me3 in the earliest replicating DNA. **(C)** B6 PRDM9 binding sites are depleted in early replicating DNA (measured by Affinity-Seq) in the B6^wt/wt^ genome. This is indicative of hotspot erosion. **(E)** Inter-homolog repair products (crossovers + non-crossovers; (Li et al., 2019a)) are depleted in the earliest replicating DNA relative to DSB-associated H3K4m3 (Note that H3K4m3 are the same data as in D, plotted against RT in B6xCAST). Simulated RT from the T1 Meiocyte population (Figure 3D) is used for B,C and D. Simulated RT from the Meiotic S-phase in B6xCAST F1 mice is used for E. Solid lines depict the LOESS smoothed signal ± standard error (shaded). For all panels, dots represent the average signal from all autosomal bins in the genome for each RT quantile (N_bins_ = 250).

We found that all measures of recombination are enriched in the early replicating regions (ERRs) (Figure 4). Nonetheless, informative differences are apparent. PRDM9-mediated H3K4m3 and DSB frequency (SPO11-oligo) are similarly enriched relative to RT (Figure 4B; yellow and magenta lines). Both profiles “flatten” in the earliest replicating DNA. This likely reflects hotspot erosion (Boulton et al., 1997; Myers et al., 2010), a process by which strong PRDM9 binding sites are purged from the genome (Figure 4C). Hotspot erosion is indicative of ancestral recombination, reinforcing the strong association between RT and recombination in mice. These findings also imply that the action of PRDM9 decouples replication and recombination through the erosion of PRDM9 binding sites. In contrast to PRDM9-mediated H3K4me3 and SPO11-oligo density, DSB repair intermediates (DMC1-SSDS, RPA-SSDS) are relatively depleted in the very earliest replicating regions (Figure 4B,D, S15). SSDS captures both the frequency (Lange et al., 2016) and the lifetime (Pratto et al., 2014) of DSB repair intermediates. Since DSB frequency closely mirrors PRDM9-mediated H3K4me3, we conclude that DSBs forming in early replicating regions are more rapidly repaired than those elsewhere. In agreement with this hypothesis, the signature of rapid repair is no longer seen in DMC1-SSDS from a mouse in which all meiotic DSBs remain unrepaired (*Hop2^-/-^*; Figure S15). The signature of rapid DSB repair in ERRs is also seen in mice that lack PRDM9 (*Prdm9^-/-^*; Figure S15). Thus, this phenomenon is independent of the mechanisms that determine the local patterning of meiotic DSBs.

Rapid DSB repair in meiosis has been proposed as a hallmark of crossover-biased DSB repair (Brick et al., 2018a; Hinch et al., 2019). To simplify the study of repair outcomes we turned to a mouse homozygous for a “humanized” PRDM9 allele; this allele of PRDM9 has a binding preference not found naturally in mice, and therefore, has left no footprint of hotspot erosion in the mouse genome. Indeed, in these mice, PRDM9-mediated H3K4me3 is linearly correlated with RT in the earliest replicating DNA (Figure 4D; yellow line), while the DMC1-SSDS signal still shows a relative depletion indicative of rapid repair. We found that all inter-homolog repair products (COs and NCOs) were notably depleted in the earliest replicating DNA, where rapid DSB repair is occurring (Figure 4E, S15). The “missing” repair outcomes likely result from DSBs that use the sister chromatid as a repair template, as inter-sister repair products cannot be detected (sister chromatids are genetically identical). Inter-sister DSB repair is generally disfavored in meiosis to assure crossover formation between homologs. Nonetheless, this inter-homolog bias appears to be gradually established, such that the earliest-forming DSBs can still repair from the sister chromatid (Joshi et al., 2015; Sandhu et al., 2020). Given that we find evidence for inter-sister repair of DSBs and given the fact that this type of repair is only favored at the earliest-forming DSBs in meiosis, the implication of our findings is that DSBs at the earliest replicating DNA are also formed early in meiosis. Other correlates of DNA replication, such as GC-content or chromatin structure lack any mechanism that would link them to early-forming DSBs. Therefore, the most parsimonious conclusion is that there exists a direct link between replication and recombination in male mouse meiosis.

### Chromosome-scale regulation of recombination is coupled to DNA replication

Chromosome-scale regulation of recombination is a PRDM9-independent aspect of recombination patterning. For example, short chromosomes, by virtue of their size, are statistically less likely to receive a crossover-competent DSB, yet mechanisms exist to assure that they do (Duret and Galtier, 2009; Kaback et al., 1992; Murakami et al., 2020). In mouse males, we find that short chromosomes have a higher crossover density (Figure 5A), however much of the variance in crossover density remains unexplained. We hypothesized that DNA replication may play a role because origin density varies greatly among mouse chromosomes; for example, origin density on chromosome 13 (120 Mb; 327 origins) is three times lower than on chromosome 11 (122 Mb; 1,023 origins). Indeed, we found that origin density is positively correlated with crossover density (Figure S16). Origin density is a crude estimate of replication activity that does not account for stochastic origin firing or competition between chromosomes for limiting firing factors. We therefore derived a more integrated metric of replication from RT-Sim models that measures how quickly chromosomes are replicated relative to each other (replication speed; Figure 5B, S16). Alone, replication speed predicts crossover density substantially better than origin density (Figure 5C, S16) and when coupled with chromosome size in a multiple linear regression model can explain 90% of the per-chromosome variance in crossover density (Figure 5D, S16). Strikingly, this implies that the recombination potential of chromosomes is mostly established before DSB formation.

In yeast, compensation for chromosome size is achieved by increasing the DSB density on short chromosomes (Murakami et al., 2020), but such an increase is not so apparent in mice (Lange et al., 2016). Whether differences in DNA replication modulate DSB formation or alter DSB repair outcomes remains to be explored.

### Distinct sub-telomeric patterning of DNA replication in the human male germline

In human males, like in mice, meiotic DSBs exhibit megabase-scale correlations across the genome (Figure 6A). An overt manifestation of this phenotype is seen at the ends of human chromosomes, where both DSBs (Pratto et al., 2014) and crossovers (Coop et al., 2008) are grossly enriched in males, independent of PRDM9 genotype (Figure 6B).

**Figure 6.**
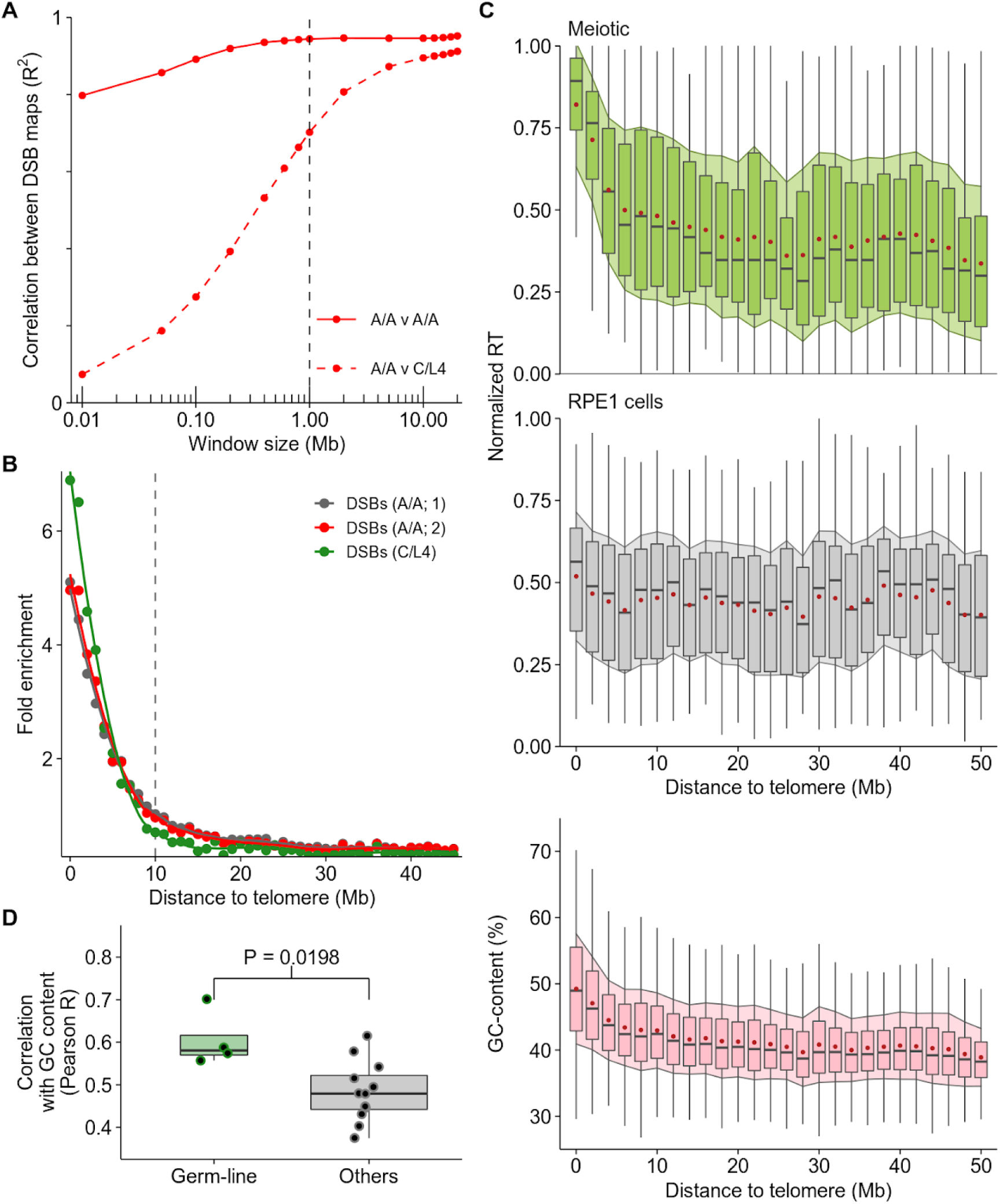
Sub-telomeric DNA replicates early in human male meiosis. **(A)** Extensive large-scale correlation in DSB density in human males with different PRDM9 genotypes (PRDM9^A/A^ homozygotes (A/A) and PRDM9^C/L4^ (C/L4) heterozygote). At fine scale (10 Kb), there is no correlation between DSB maps in the A/A and C/L4 individuals because DSB hotspot locations differ. Comparing the maps at lower resolution reveals extensive correlation at megabase scales. **(B)** Meiotic DSBs are enriched in sub-telomeric regions in human males, independent of the *Prdm9* genotype. **(C)** Subtelomeric DNA replicates consistently early in human male meiosis but not in mitotic (RPE-1; Table S3) cells. Boxplots depict the range of replication timing values in 2 Mb regions across the genome; the median is shown as a grey bar and the box designates the interquartile range. The mean (red dot) ± 1 standard deviation (filled shadow) is also shown for each interval. GC content is also elevated in sub-telomeric DNA. **(D)** RT in the germline (all meiotic samples) correlates better with genomic GC content than RT in other cell types.

We isolated S-phase nuclei expressing meiotic markers from human testes (Figure S17) and inferred replication timing. Strikingly, we found that sub-telomeric regions of human chromosomes replicate early and that therefore, germline RT is highly correlated with DSB hotspot density (R^2^ = 0.75) (Figure 6B-C). This germline-specific RT patterning may explain why only a weak link was previously found between RT in a lymphoblastoid cell line (LCL) and meiotic recombination (Koren et al., 2012). The distal pattern of early replication in male meiosis appears highly distinct, as sub-telomeric DNA does not replicate notably early in any of the other cell types we studied (Figure 6C, S18). Intriguingly, GC-content is also elevated in sub-telomeric DNA (Figure 6C,D); thus, genomic GC-content is better correlated with RT in the germline than in other cell types (Figure 6D). Importantly, the earliest meiotic DSBs in human males are detected almost exclusively in sub-telomeric DNA (Pratto et al., 2014). Thus, early DNA replication in distal regions in the germline underlies the spatio-temporal patterning of meiotic recombination in human males.

## Discussion

The single round of DNA replication that begins meiosis has remained an enigma, despite huge advances in our ability to interrogate nuclear processes. Aside from a cadre of classical papers in the 1970s, few studies have addressed the intricacies of meiotic DNA replication in mammals at a molecular level. We have addressed this shortcoming by developing a comprehensive framework for the study of DNA replication *in vivo* and have used this to generate the first major advances in the study of DNA replication in mouse and human meiosis in decades.

A major hurdle to overcome was to map origins of replication in the few S-phase cells that are found in the mammalian testis. Existing protocols to map origins are designed for the controlled environment of cell culture, typically require tens to hundreds of millions of cells, and have not been performed in tissue. By adapting methods for nascent-strand sequencing to retain the directionality of captured strands, we generated a high-confidence map of ∼11,500 origins of replication in the testis. This is also the first map of *in-vivo* replication origins in a mammal and paves the way for future studies in other cell types. Replication origins appear to be determined by the combination of open chromatin and CpG density and are unevenly distributed in the genome. CpG islands in open chromatin are broadly conserved across cell types, and indeed, we find that the origins identified in testis represent a core set of cell-type-agnostic replication origins. This does not negate the possibility of cell-type specific origins, as have been reported in other studies (Smith et al., 2016).

Despite using the same population of replication origins, we exposed fundamental differences in S-phase duration among cell-types through *in-silico* modelling. Importantly, experimental methods to infer S-phase duration are laborious and imprecise or are limited to cell culture systems (Grant et al., 2018). Meiotic S-phase has been suggested to be notably long in a variety of organisms, including animals, plants, and yeast (Bennett et al., 1972; Callan, 1973; Cha et al., 2000; Holm, 1977). Experimental estimates of S-phase duration in mice are 14-15 hr and 29-30 hr in Spermatogonia-B and Spermatocytes, respectively (Ghosal and Mukherjee, 1971; Monesi, 1962). In agreement, we found that meiotic S-phase in mice is approximately 1.8 times longer than the germ-cells that immediately precede meiosis. The similarity between *in-silico* and experimental estimates implies that the assumptions of our model are reasonable; this includes the assumption that replication fork speed is similar across cell-types. To accurately model replication, a limiting factor that caps the number of active replisomes is strictly required ((Gindin et al., 2014) & data not shown). We found that this replisome “ceiling” differs among cell types, and, in turn, modulates S-phase duration. Importantly, such estimates of the number of active replisomes per cell are extremely challenging to obtain experimentally (Chagin et al., 2016). What components of the replication machinery govern this “ceiling” remain unknown, but investigating the factors that slow down meiotic S-phase may help identify genes that modulate S-phase duration more generally. In yeast, the meiosis-specific cohesin Rec8p and DSB-forming protein Spo11p appear to play such a role. However, we found no evidence for a role of SPO11 in regulating S-phase duration in mouse meiosis. In addition to a lowered replisome “ceiling”, other, more complex mechanisms may contribute to slowing S-phase in meiosis. Our model only allows alteration of the global properties of replication, however regional modifiers of replication fork speed such as replication slow zones (Cha and Kleckner, 2002) or common fragile sites (Smith et al., 2006) may impede fork progression differentially in meiosis and other cell types.

Our tripartite approach to describe and parameterize DNA replication enabled us to ask if replication influences recombination in mammalian meiosis. We found an extremely strong positive correlation between multiple metrics of recombination and early replicating DNA. The earliest measure we examined was the binding of the PRDM9 protein, where we found that the preference for DSB formation in early replicating DNA was already established. One possibility therefore is that opportunistic binding of PRDM9 to more accessible chromatin in early replicated DNA establishes a link between replication and recombination. However, we found that recombination was enriched in early replicating DNA even in the absence of functional PRDM9. This implies that broad-scale patterning of recombination acts independent of the factors that determine the local patterning of meiotic DSBs.

We found strong evidence for the rapid repair of meiotic DSBs in early replicating parts of the genome. The rapid repair of DSBs is a correlate of crossover-biased resolution (Hinch et al., 2019), however, we find that all interhomolog repair outcomes are depleted in these regions suggesting increased use of the sister-chromatid as a repair template. In meiosis, DSB repair with the sister chromatid is strongly disfavored. This assures that recombination occurs with the homologous chromosome - a requirement for allelic exchange and accurate chromosome segregation. At least in yeast, this inter-homolog bias is established gradually, and inter-sister recombination is preferred only for the earliest DSBs that form in meiotic prophase (Joshi et al., 2015; Sandhu et al., 2020). We therefore infer that the earliest DSBs occur in the earliest replicating DNA, strongly implicating a mechanistic link between replication timing *per se* and DSB formation. This would also imply that correlates of replication timing such as GC-content, genome compartmentalization, heterochromatin, or gene density, which lack any temporal component, may indirectly affect recombination by modifying replication patterns. In human males, cytological evidence has shown that the earliest meiotic DSBs occur almost exclusively in sub-telomeric DNA (Pratto et al., 2014). This coincides with the earliest replicating DNA, substantiating the direct link between replication and recombination initiation. It remains possible that there are two phenomena that modulate recombination at large-scales in the genome; one that temporally favors early-forming DSBs in the wake of a passing replication fork (like in yeast), and another that favors recombination in the resultant “permissive” chromatin environment. Of course, since DNA replication establishes the 3D structure of the genome (Klein et al., 2019), these two effects may be one-and-the-same.

It is well established that chromosome size partly determines per chromosome recombination rates (Duret and Galtier, 2009; Kaback et al., 1992; Murakami et al., 2020). In addition, we found that in mouse males, the per-chromosomes origin density has a similar predictive value to chromosome size. Importantly, the predictive power of chromosome size and replication are additive, which implies that these properties represent uncorrelated aspects of the mechanisms that control per chromosome recombination rates. Together almost all of the per chromosome variation in crossover density can be explained by these two properties, suggesting that recombination will follow a deterministic path, established before DSBs are made. Short chromosomes likely benefit from an elevated recombination rate to assure crossover formation, however it is less intuitive to understand if or why elevated recombination on fast replicating chromosomes would be beneficial. Fast-replicating chromosomes are origin rich, GC-rich, replicate earlier, and have elevated gene density. A mechanism that links replication to recombination would assure that such regions benefit from recombination to break linkage blocks, generate diversity and purge deleterious mutations.

Any benefit of recombining in early replicating DNA could only manifest in the germline (where meiotic recombination occurs) and intriguingly, in human males, germline DNA replication mirrors GC content more closely than replication in other cell types. Thus, during development, we propose that replication may follow a well-charted course, dictated by the underlying DNA. In contrast, the commitment to differentiation in other cell types may render replication more susceptible to epigenetic regulation (Hiratani et al., 2010). In being closely linked to GC content, DNA replication may act as a “selectable trait” that influences the evolution of the genome. For example, should a chromosome lack sufficient recombination capacity in meiosis, there would be a strong selective advantage to increasing GC-content, leading to earlier replication and hence, elevated DSB potential.

Errors in DNA replication, if unrepaired, can alter the genome, but will only be transmitted to the next generation if they occur in the germline. Our finding that germline replication is substantially different therefore has important implications for understanding the patterning of *de-novo* variation in the genome and its impact on population genetic structure.

## Acknowledgments

We acknowledge all members of the Camerini-Otero lab for helpful discussion and comments. We also thank Galina Petukhova, Michael Lichten and Peggy Hsieh for critiquing the manuscript. We thank the Flow Cytometry Core of NHLBI, the NIDDK Laboratory of Animal Sciences Section and the NIDDK genomics core facility. This work utilized the computational resources of the NIH HPC Biowulf cluster (http://hpc.nih.gov). This work was funded by an NIGMS grant to P.W.J. (R01GM11755) and by the intramural program of the NIDDK (R.D.C.O.). We also thank the Washington Regional Transplant Community for their assistance in obtaining de-identified human testis donations for research.

## Author Contributions

Conceptualization, F.P., K.B. and R.D.C.O.; Methodology, F.P., K.B., G.L. and G.C.; Investigation, F.P., K.B., G.L., D.D., J.C. and G.C.; Formal Analysis, F.P., K.B.; Software, K.B.; Visualization, F.P., K.B.; Writing - Original Draft, F.P. and K.B.; Writing - Review & Editing, F.P., K.B. and R.D.C.O.; Resources, S.R.W. and P.W.J.; Funding Acquisition, P.W.J. and R.D.C.O.; Supervision, R.D.C.O.

## Declaration of Interests

The authors declare no competing interests.

## Data & code availability

The sequencing data have been deposited at the GEO (accession number GSE148327, reviewer token XXXXXXX). Custom pipelines and code are available at github:

https://github.com/kevbrick/prattoEtAlAnalyticPipeline.git

https://github.com/kevbrick/RTSeqPipeline.git

https://github.com/kevbrick/RTsim.git

## Supplementary Figures

**Figure S1.**
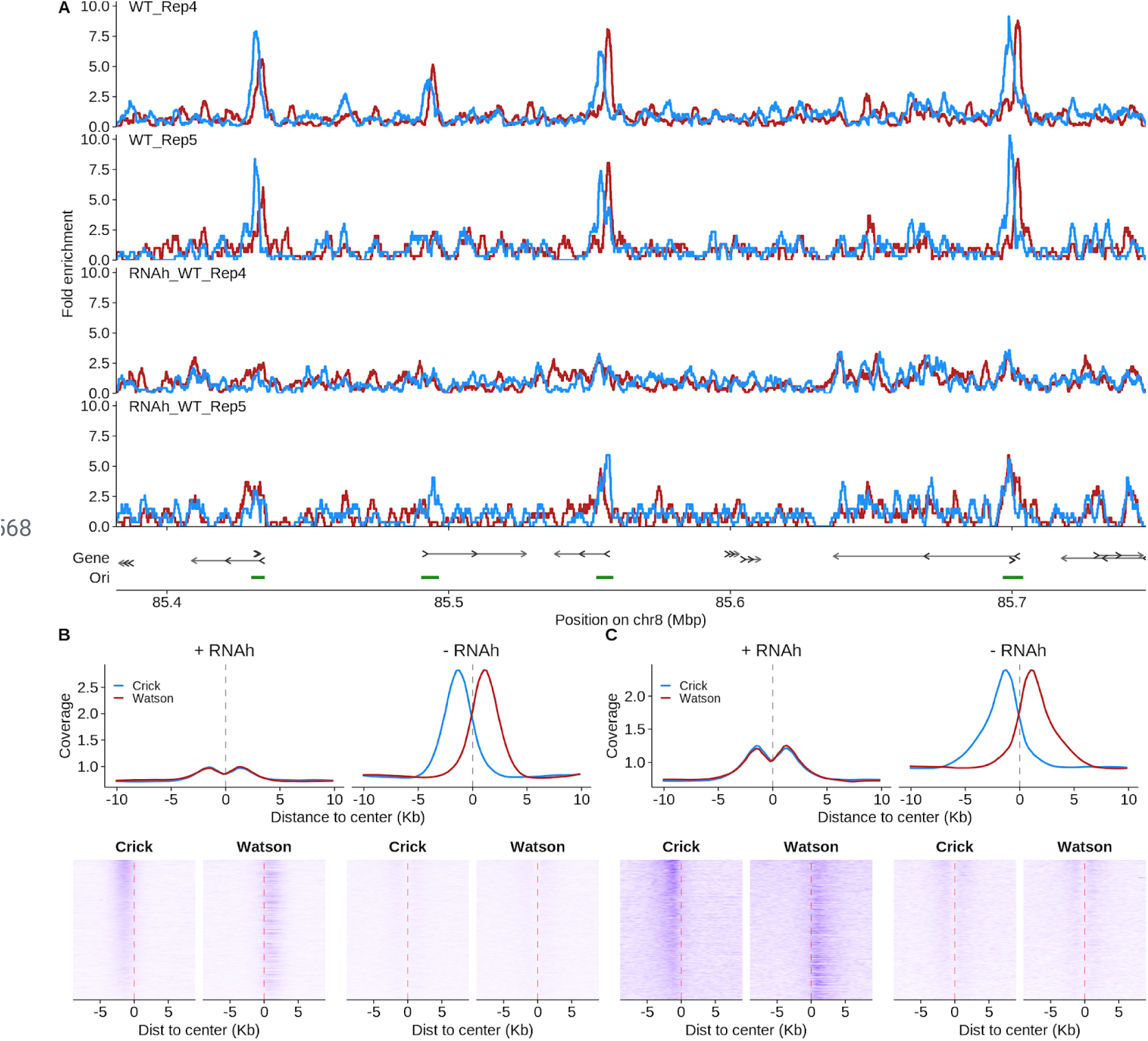
RNA hydrolysis abolishes the Ori-SSDS signal. RNA hydrolysis prior to lambda exonuclease treatment abolishes Watson/Crick asymmetry at origins of replication. This is because RNA hydrolysis will remove the nascent strand RNA primer and thus render nascent-strands susceptible to lambda exonuclease digestion (see methods). Two replicate experiments were performed in parallel; WT_Rep4 & WT_Rep5 (see methods) **(A,B,C)** The strand asymmetry of Ori-SSDS is lost following RNA hydrolysis (b,c RNAh+ samples). Residual enrichment at origins, without Watson/Crick asymmetry is still seen. This is likely because there is more DNA in the population at these sites as a result of DNA replication. Upper panels in B,C depict the average Ori-SSDS signal at origins of replication for Watson (red) and Crick (blue) strand reads.

**Figure S2.**
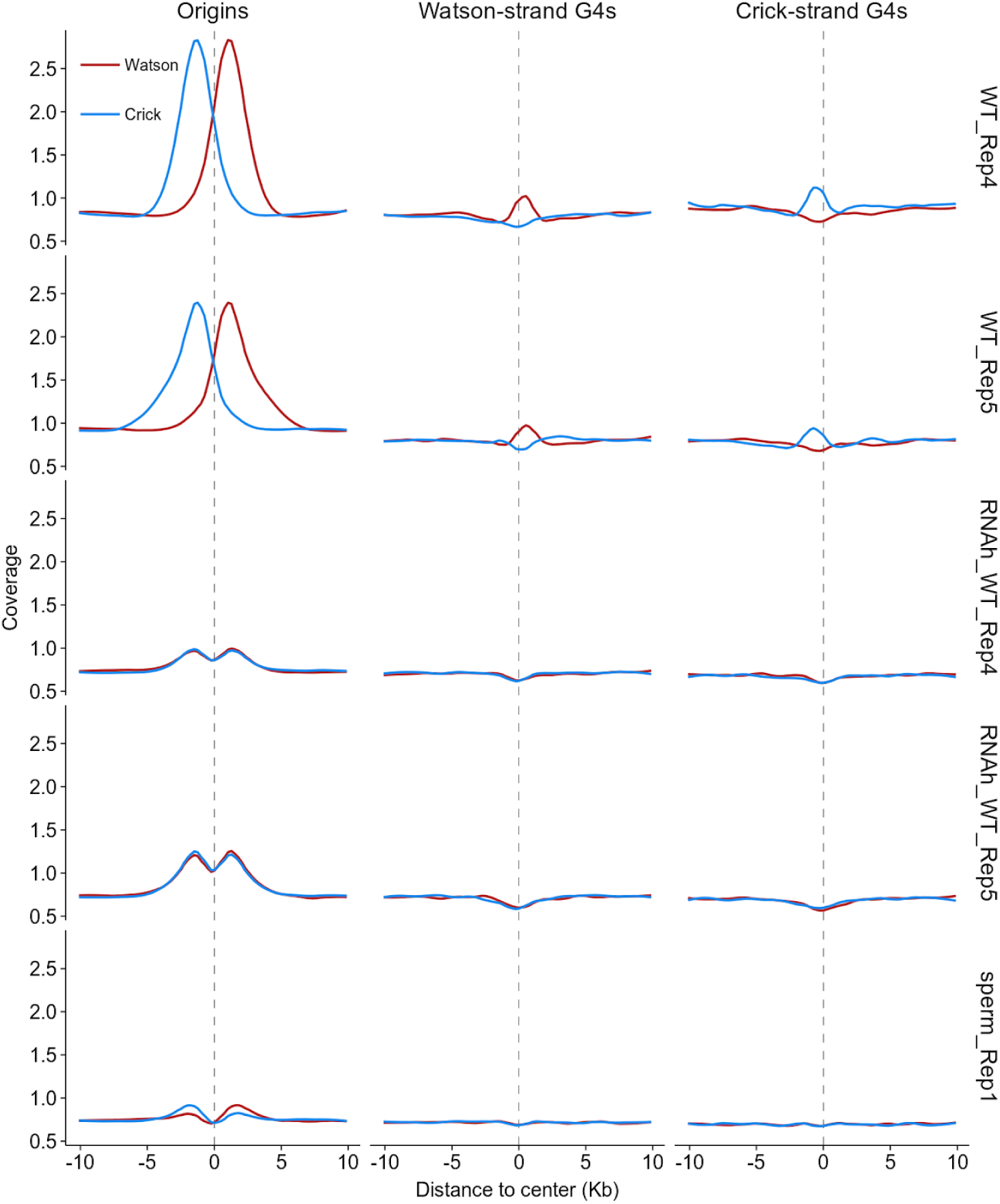
Ori-SSDS signal at putative g-quadruplexes in replicating cells. G-quadruplexes (G4s) are secondary structures that have been hypothesized to impede lambda exonuclease *in vitro* and may result in SNS-Seq peak artifacts (Foulk et al., 2015). Indeed, we detect small peaks at putative G4 sequences in Ori-SSDS data. Figures depict the average Ori-SSDS signal at origins of replication or predicted G-quadruplexes for Watson (red) and Crick (blue) strand reads. Ori-SSDS sequencing can distinguish G4-signals from origin-derived signals because G4s yield ssDNA on just one strand whereas reciprocal Watson/Crick asymmetry is seen around the center of origins. This differentiates Ori-SSDS from other SNS-Seq methods, which cannot distinguish G4s from origins. At origins that coincide with putative G4s (Huppert and Balasubramanian, 2005) it is not possible to resolve these signals however, since the G4 signal is far weaker than that at origins, the contribution of G4s to true origin signal is a minor concern. Curiously, the signal at G4s is not seen in non-replicating tissue (sperm) or in samples where the RNA-primer is degraded. Thus, the G4-associated signal is replication-dependent and not simply the result of G4 resistance to lambda exonuclease. The stranded-ness of the signal is consistent with the capture of ligated Okazaki-fragments at the G4 sites. This implies that G-quadruplexes are an impediment to fork passage *in-vivo*.

**Figure S3.**
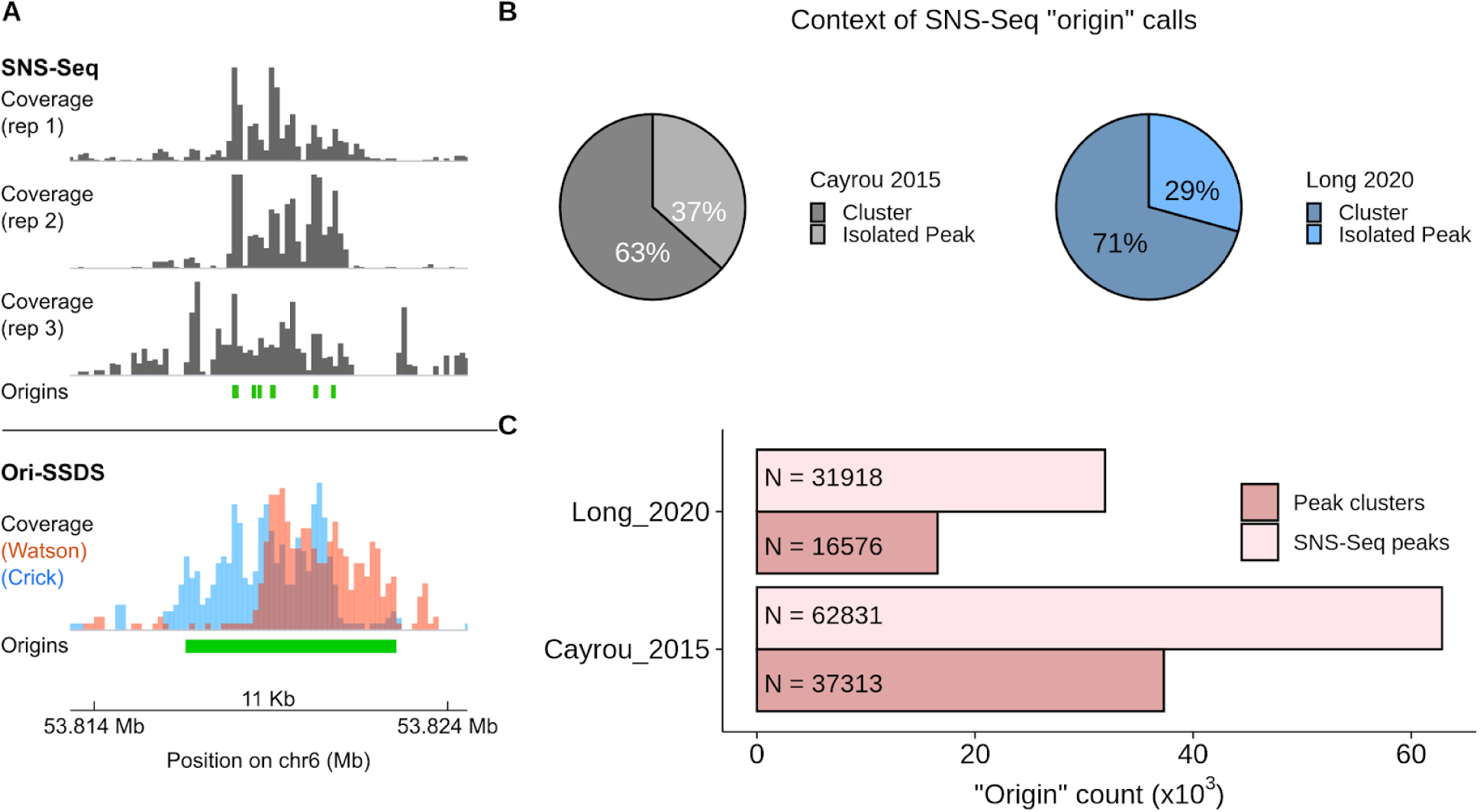
SNS-Seq peaks occur in clusters at individual origins of replication. **(A)** Without knowledge of strand-asymmetry, SNS-Seq experiments have historically overestimated the number of origins in the genome. At this locus, Ori-SSDS reveals that 6 SNS-Seq origin calls likely represent a single origin of replication. Grey tracks represent three replicate SNS-Seq coverage tracks (Cayrou et al., 2015). Green boxes underneath the grey tracks represent SNS-Seq origin calls (from GEO: GSE68347_Initiation_Sites.bedGraph.gz; accession GSE68347; UCSC liftover was used to convert mouse mm9 to mm10 genome coordinates) (Cayrou et al., 2015). Ori-SSDS coverage from WT Rep 1 is shown. The green box under the Ori-SSDS track represents the single Ori-SSDS origin call in this region. This is validated by the characteristic Watson-Crick asymmetry. **(B)** We obtained SNS-Seq origin of replication peak calls (origins) for mouse (described in panel a.; (Cayrou et al., 2015)) and for human (from GEO; GSE134988_siNC-NS_peaks.bed.gz; accession GSE134988) (Long et al., 2020). For each dataset, origins were extended ± 1.5 Kb from the center-point and merged into origin “clusters”; clusters >10 Kb were discarded. We posit that each of the remaining clusters represents a single true origin of replication. Most SNS-Seq peaks occurred in a cluster. Just 37% and 29% occurred at isolated origins in mouse and human, respectively. **(C)** The number of “origins of replication” from SNS-Seq studies is substantially reduced if clusters are considered as single origins.

**Figure S4.**
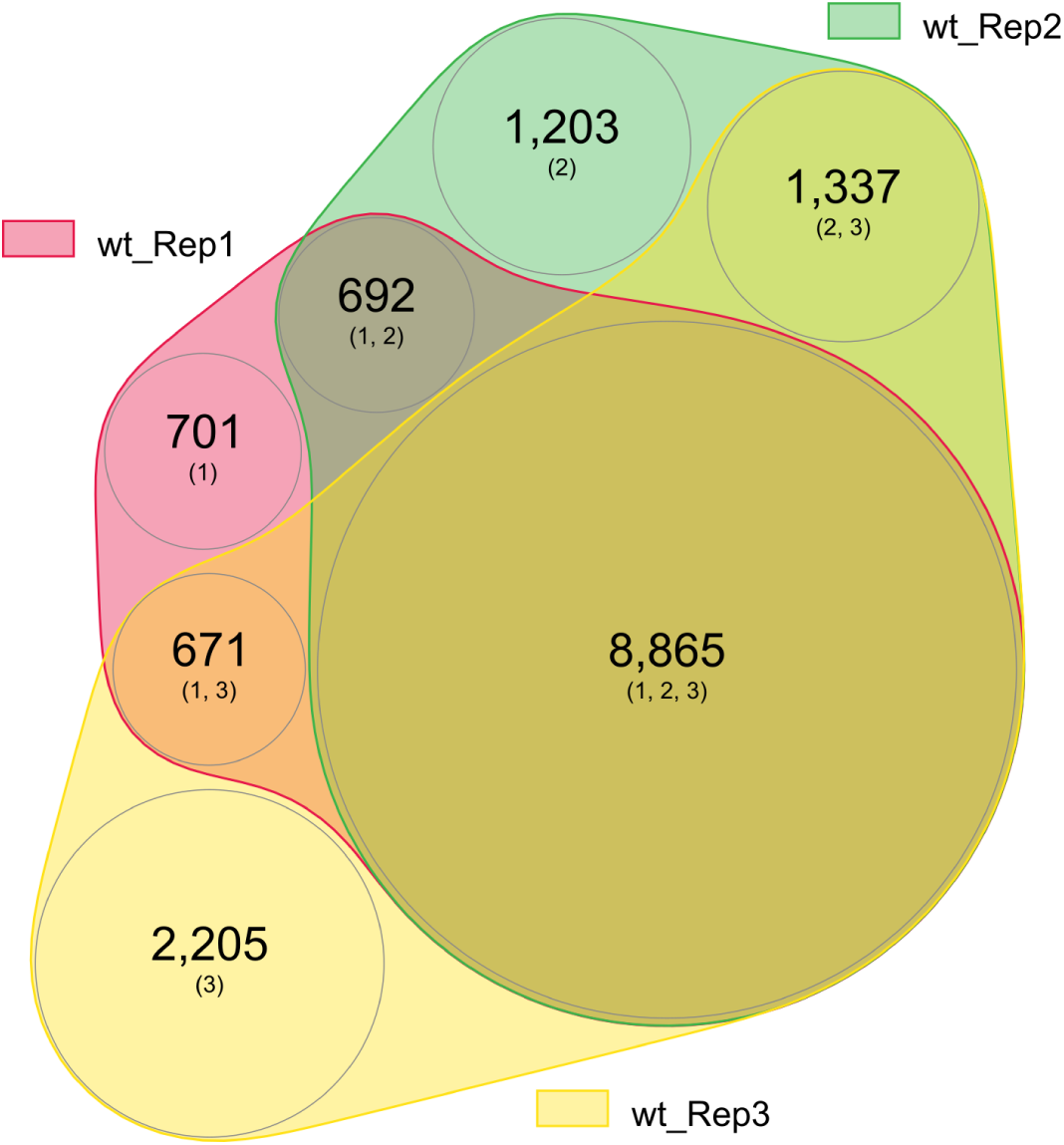
Most origins are detected in multiple replicate experiments. Proportionate venn diagram depicting the number of origins detected in each of three replicate Ori-SSDS experiments and the overlaps between sets. Our consensus set of origins includes all origins found in at least two Ori-SSDS experiments.

**Figure S5.**
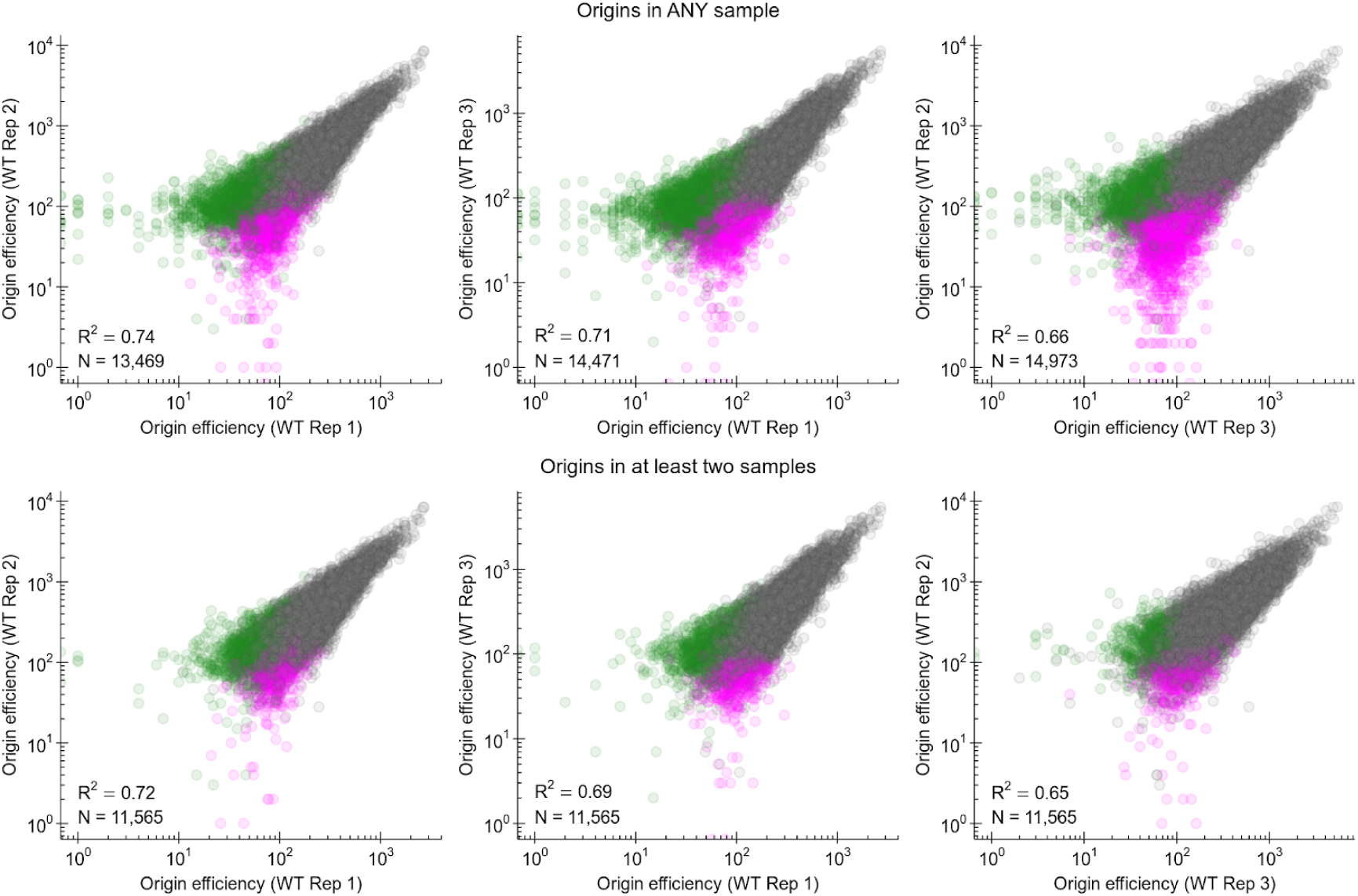
Ori-SSDS gives a reproducible readout of origin efficiency. Origin efficiency is a measure of the frequency at which each origin is used. This is inferred from the sequencing read coverage in each experiment (see methods). Grey dots represent origins found in both samples. Magenta and green dots represent origins found only in the x-axis designated and y-axis designated samples, respectively. The top panels show all origins found in either experiment for each pair-wise comparison. The bottom panels show only the consensus set of 11,565 origins. R^2^ is the squared Spearman correlation coefficient of log transformed values. N indicates the number of origins for each pairwise comparison.

**Figure S6.**
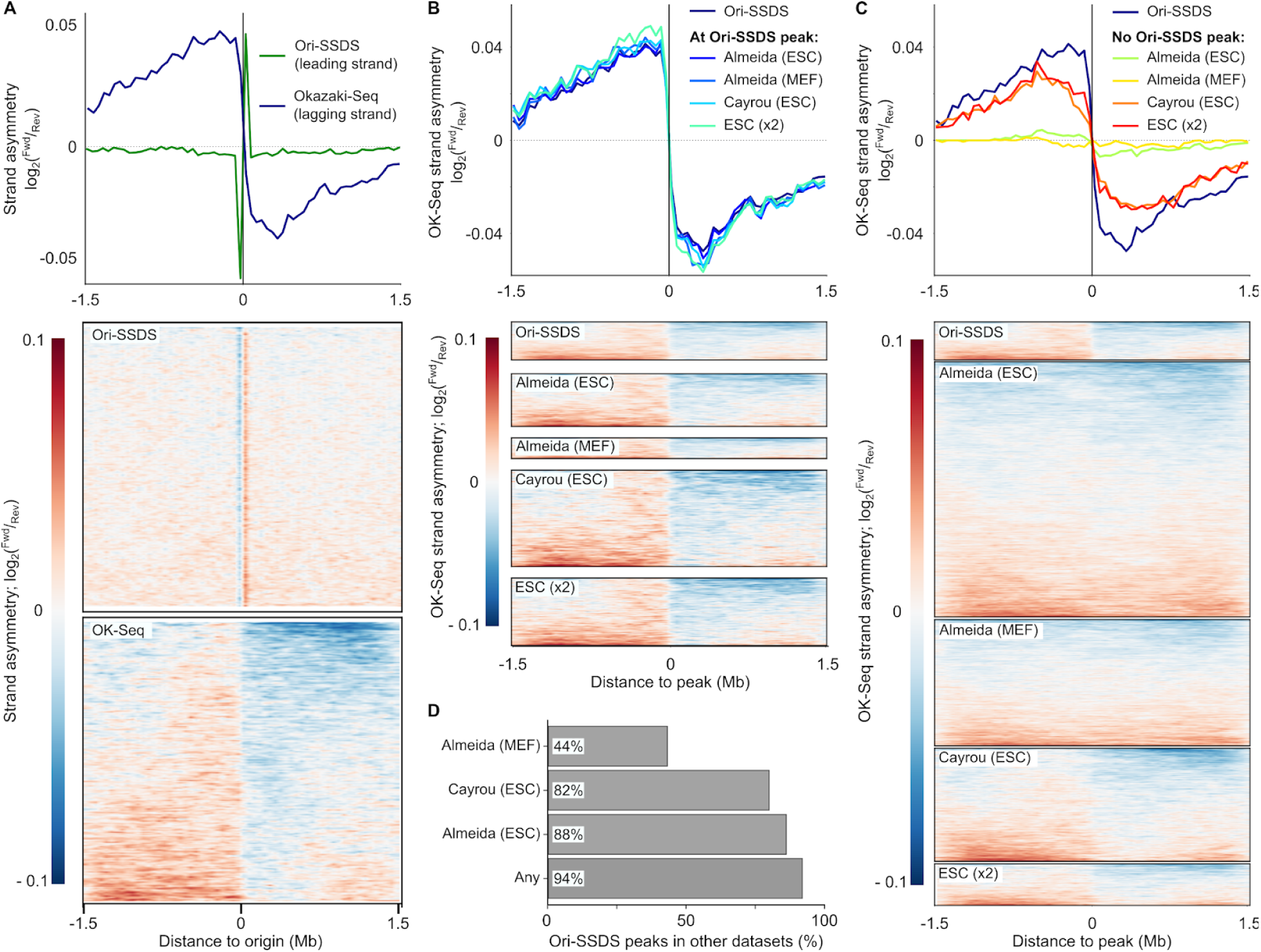
Ori-SSDS peaks coincide with replication measures from other experiments. **(A)** Okazaki fragments are distributed asymetrically around Ori-SSDS-defined origins of replication. Okazaki fragment sequencing (Ok-Seq) data in Embryonic Stem Cells (ESCs) are from (Petryk et al., 2018). The Okazaki fragment signal is far broader than the signal from Ori-SSDS because it captures all lagging-strand synthesis (for details, see (Petryk et al., 2016)); Ori-SSDS captures only RNA-primed leading strands at the origin of replication. Note that Ok-Seq exhibits the opposite polarity to the Ori-SSDS signal as expected from capture of lagging and leading strands, respectively. Data are shown in 50 Kb windows. Heatmaps depict the signal at individual Ori-SSDS defined origins. **(B,C)** Ok-Seq signal at subsets of origins of replication. For reference, the Ok-Seq signal at Ori-SSDS defined origins is shown in three panels (A,B,C). The heatmaps in panels B & C are scaled to reflect the number of origins in each subset. There are three genome-wide maps of origins of replication in mice. All three maps were generated using SNS-Seq. One map is derived from Mouse Embryonic Fibroblast (MEF) cell culture (Almeida et al., 2018) and two from ESC culture (Almeida et al., 2018; Cayrou et al., 2015). **(B)** Origins defined in SNS-Seq experiments that coincide with Ori-SSDS peaks exhibit similar Ok-Seq asymmetry. Data for origins common to the two SNS-Seq experiments in ESCs are also shown (ESC (x2)). **(C)** A large proportion of peaks detected in SNS-Seq experiments, but not in Ori-SSDS appear to be false-positives as they lack Ok-Seq asymmetry. This is more pronounced in the Almeida experiments than in the Cayrou experiment. **(D)** Most origins of replication from Ori-SSDS in testis coincide with origins mapped in other cell types. Origins from the three SNS-Seq experiments were combined to assess the overlap with “Any” origins defined previously.

**Figure S7.**
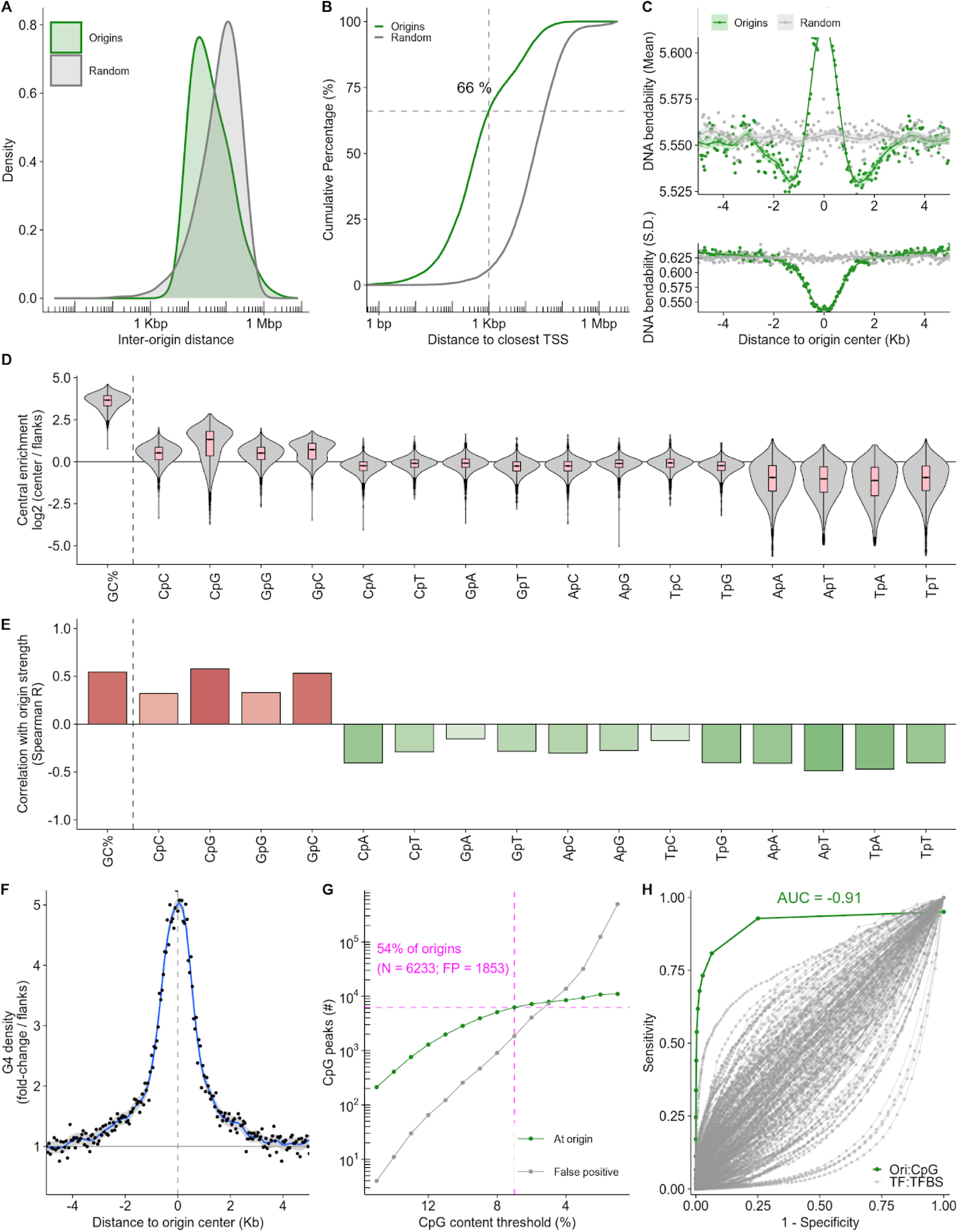
Origins of replication in mouse testis occur at GC-rich, accessible sites. **(A)** Origins of replication are more closely spaced than expected in the genome. The expected distances between randomly chosen intervals is shown in grey. **(B)** Most origins occur near gene promoters; 66% occur within 1 Kbp of a transcription start site (TSS), far more than expected (random; grey line). **(C)** Replication origins coincide with local changes in DNA bendability (calculated from (Goodsell and Dickerson, 1994)). The mean (top) and standard deviation (bottom) of the DNA bendability at all origins is shown. Each point represents a 100 bp window. Solid line is a LOESS fit. **(D)** Origin centers are GC-rich and enriched in CpG and GpC dinucleotides. The log_2_ ratio of the central 500 bp compared to the region flanking this center is shown. **(E)** GC-content (GC%), CpG density and GpC density correlate positively with origin efficiency. **(F)** Putative G4-forming sequences are highly enriched at the center of origins. 68% of origins have a putative G4-forming sequence in the central ± 1 Kb. **(G)** Origin locations can be identified from CpG density. CpG dinucleotide density was evaluated in all 1 Kb regions in the genome using a 100 bp sliding window. Intervals at each CpG threshold were expanded ± 100 bp around the center and overlapping windows were merged. The centerpoint of each merged interval was defined as a CpG peak. CpG peaks with CpG content ≥ 7% overlap 6,233 origins (53% of total). 1,853 CpG peaks at this threshold do not occur at an origin (FP). **(H)** Origins are better predicted by CpG density than transcription factors (TFs) are by transcription factor binding sites (TFBS). Receiver operating curve (ROC) for predicting origin locations using CpG peaks (green) or for predicting peaks in ChIP-Seq datasets for 179 transcription factors (grey). For origins, at each CpG threshold (1% to 15% in 1% steps), peaks were called (as in G). All mouse TFs with ChIP-Seq data (in GTRD database; (Yevshin et al., 2017)) and with a designated TFBS in the HOCOMOCO database (Kulakovskiy et al., 2018) were used. SPRY-SARUS (Kulakovskiy et al., 2018) was used to identify TFBS motifs genome-wide with p<0.005. For each TF, the score at which <= 12,000 peaks were found was used as a lower threshold (this is to assure a fairer comparison with origins). Motifs with scores less than this were discarded. TFs with <12,000 peaks not used. Hits inside ChIP-Seq peaks were considered true positives.

**Figure S8.**
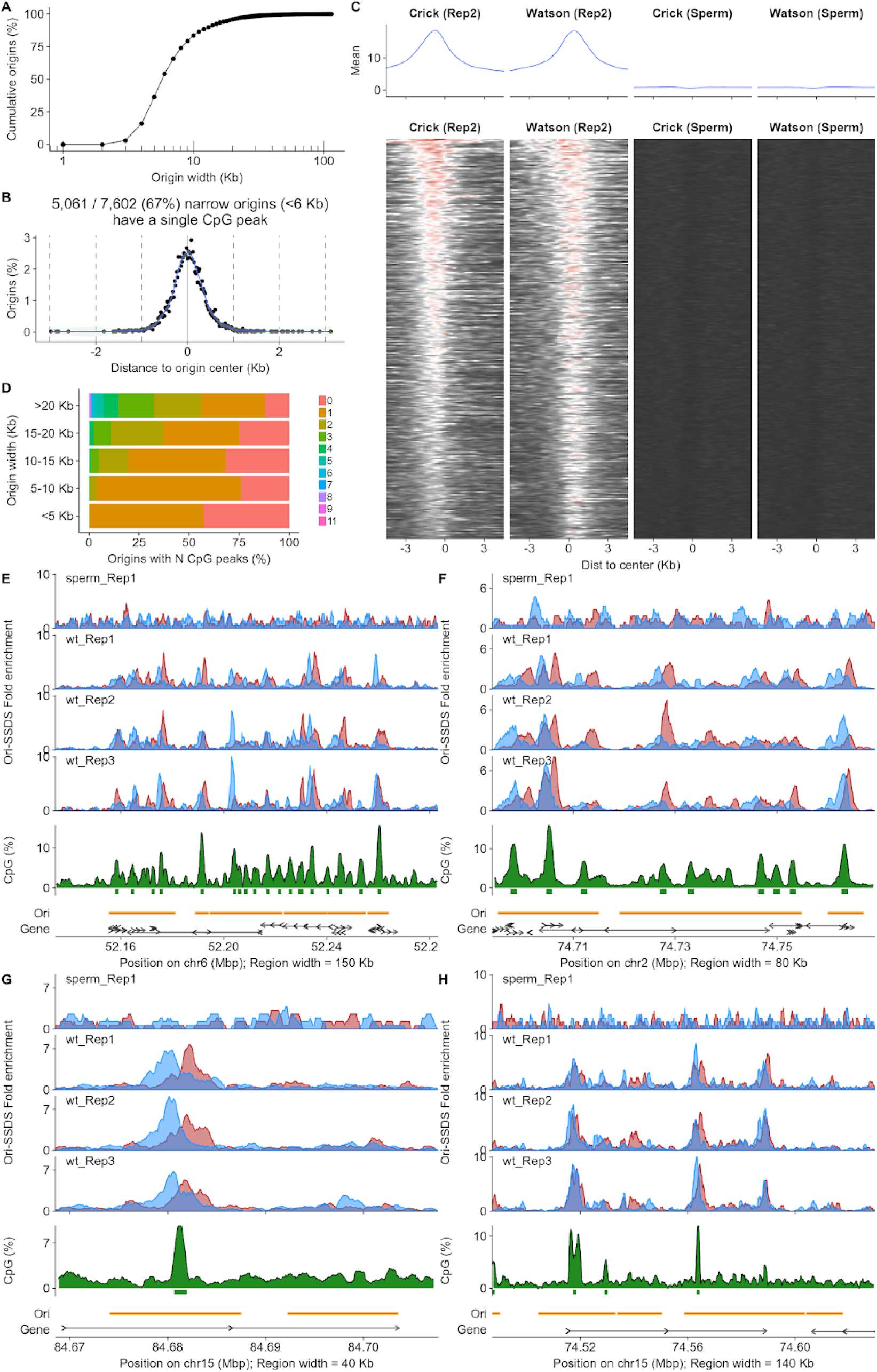
Deconvolution of closely spaced origins of replication. **(A)** Origin of replication cumulative width distribution. **(B)** Peaks of CpG density coincide with origin centers. Only origins narrower than 6 Kb were considered. CpG dinucleotide density was evaluated in all 1 Kb regions in the genome using a 100 bp sliding window. Intervals with >5% CpGs were expanded ± 100 bp around the center and overlapping windows were merged. The centerpoint of each merged interval was defined as a CpG “peak”. **(C)** Almost all CpG peaks within broad initiation zones (origins >10 Kb) exhibit Ori-SSDS Crick-Watson asymmetry. This strongly implies that these are individual origins of replication. **(D)** Local CpG peak counts as a function of origin width. **(E,F)** Examples of a cluster of initiation zones. Peaks of CpG density coincide with Crick/Watson asymmetry at many loci, revealing extensive sub-structure. This locus likely contains many discrete origins of replication that are difficult to separate *in silico*. The Hox locus is shown in (E). **(G)** Two adjacent initiation zones with different properties. The zone on the left is 13 Kb wide, yet contains a single, clear CpG peak that coincides with Ori-SSDS asymmetry. This is likely a single replication origin. The zone on the right is also 13 Kb, but contains no discernable CpG peak or origin. This may represent a diffuse initiation zone or a cluster of weak origins. **(H)** Four initiation zones with varying properties.

**Figure S9.**
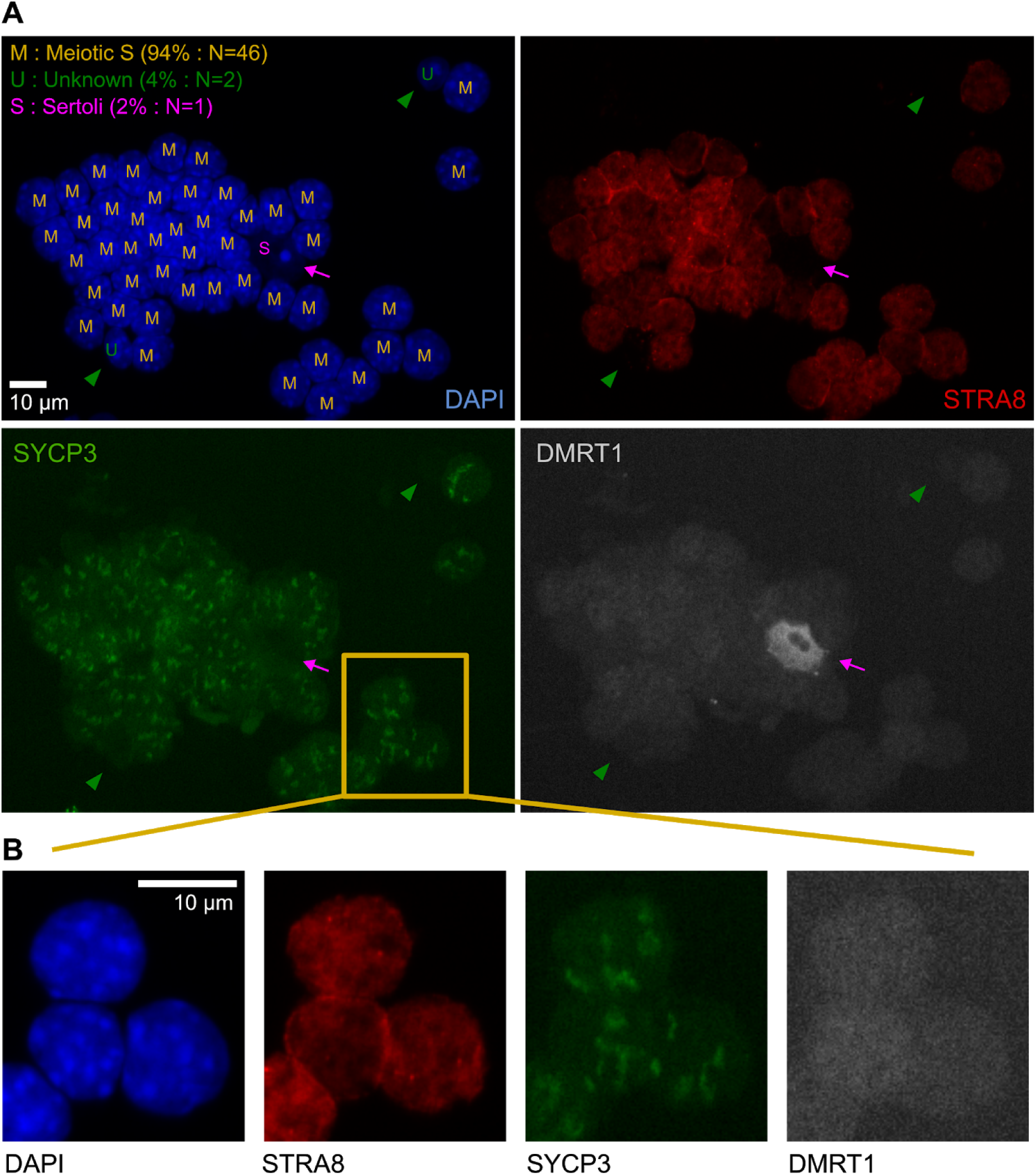
High purity populations of sorted meiotic S-phase nuclei. **(A)** Immunofluorescence microscopy images of sorted nuclei from the MeiS population (same population as in Fig. 2C; 40X magnification). Nuclei were concentrated in a small volume after centrifugation. The nuclei suspension was then pipetted onto silane coated slides and mounted with Vectashield. 49 nuclei are shown of which 46 are preleptotene nuclei (STRA8 positive, DMRT1 negative, weak and diffuse SYCP3 signal and DAPI morphology consistent with preleptotene cells (Bellve et al., 1977)). The magenta arrow points to a Sertoli cell nucleus and green arrowheads point to nuclei of unknown type. Nucleus types are indicated by a letter code superimposed on each cell of the DAPI-stained image (top-left). **(B)** Details of three preleptotene nuclei at higher magnification.

**Figure S10.**
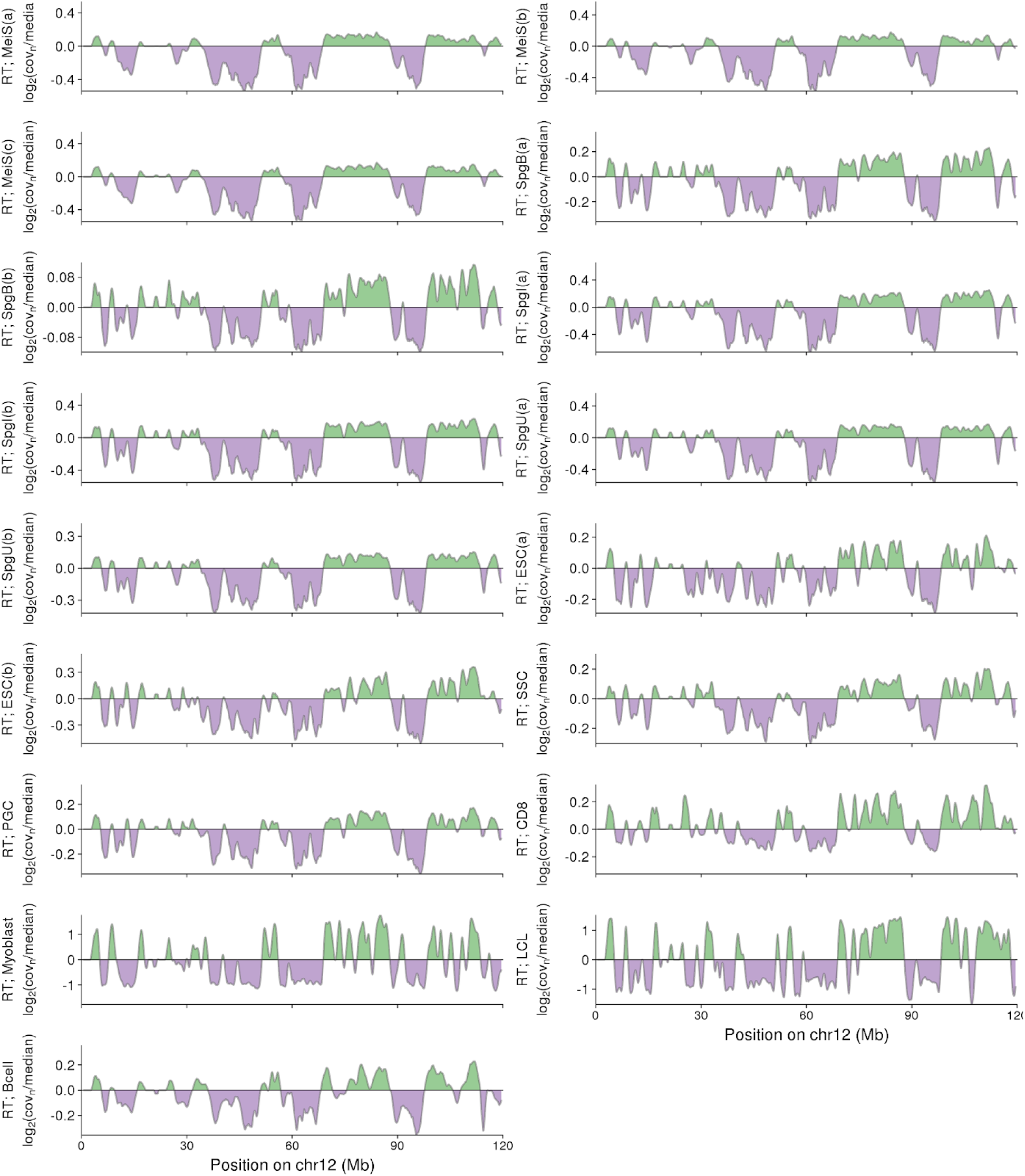
RT-Seq yields similar RT profiles across varied cell-types. RT for all cell types shown on Figure 2. RT is expressed as the log_2_ enrichment over genomic median coverage for all panels except for Myoblast and LCL data. These data were obtained in processed form from the Replication Domain database. All panels depict chromosome 12. Sample names are explained briefly in Figure 3 and in detail in Tables S1-3.

**Figure S11.**
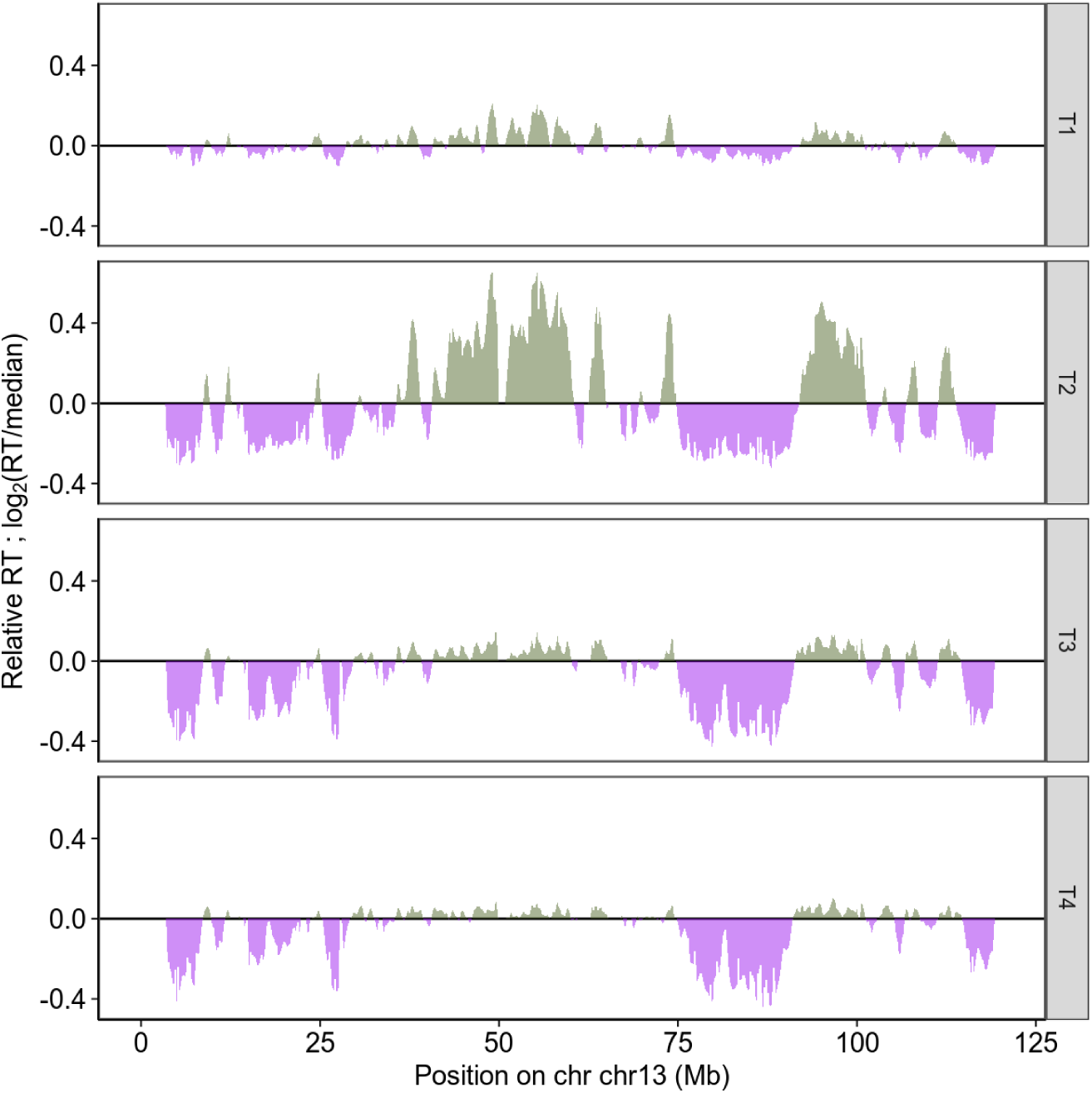
Experimentally derived RT varies as a function of the starting population. Four sub-populations of meiotic S-phase were isolated from early S (T1) through late S (T4). RT was inferred from WGS (see methods). The earliest and latest replicating regions exhibit substantial differences in RT across these four populations.

**Figure S12.**
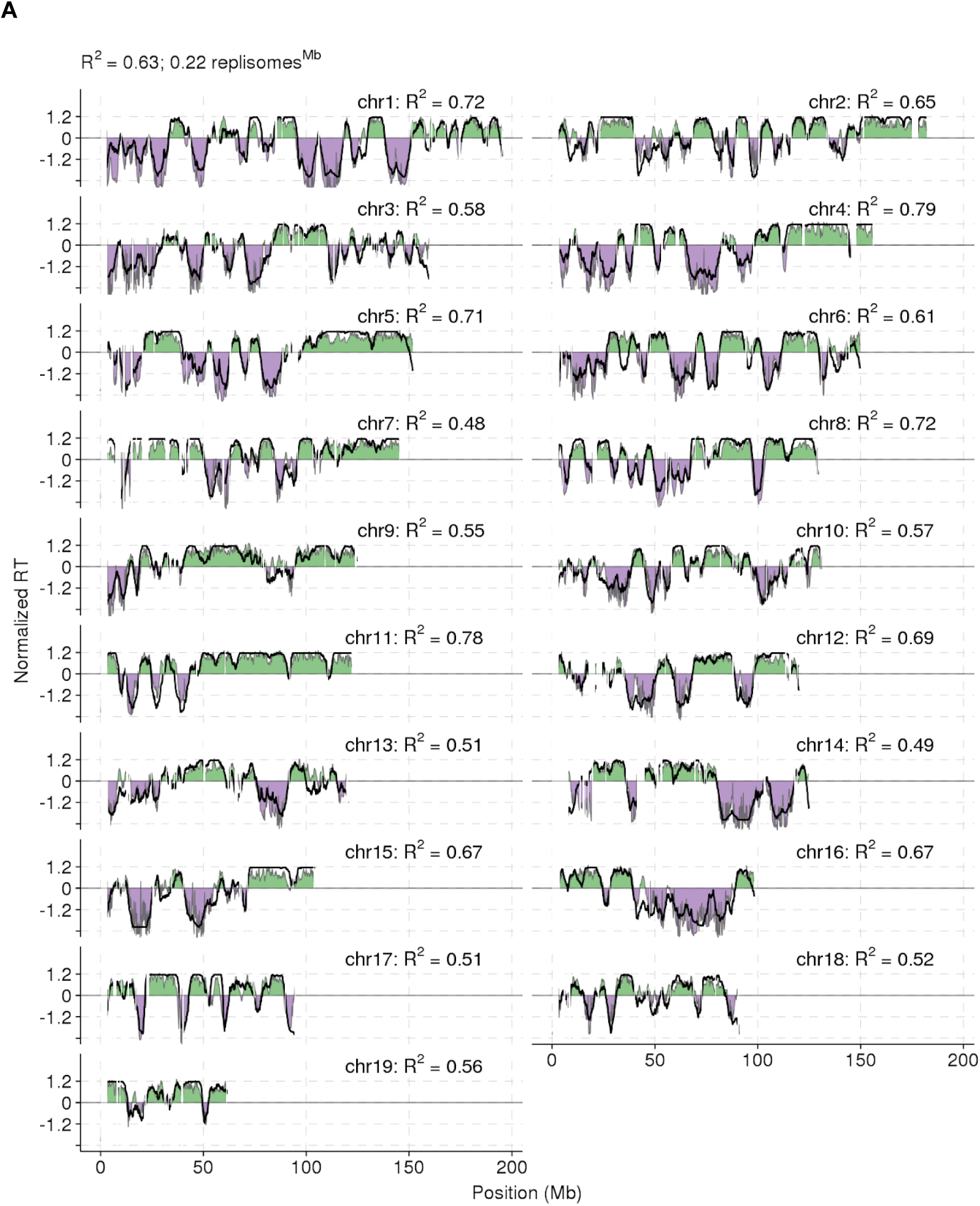
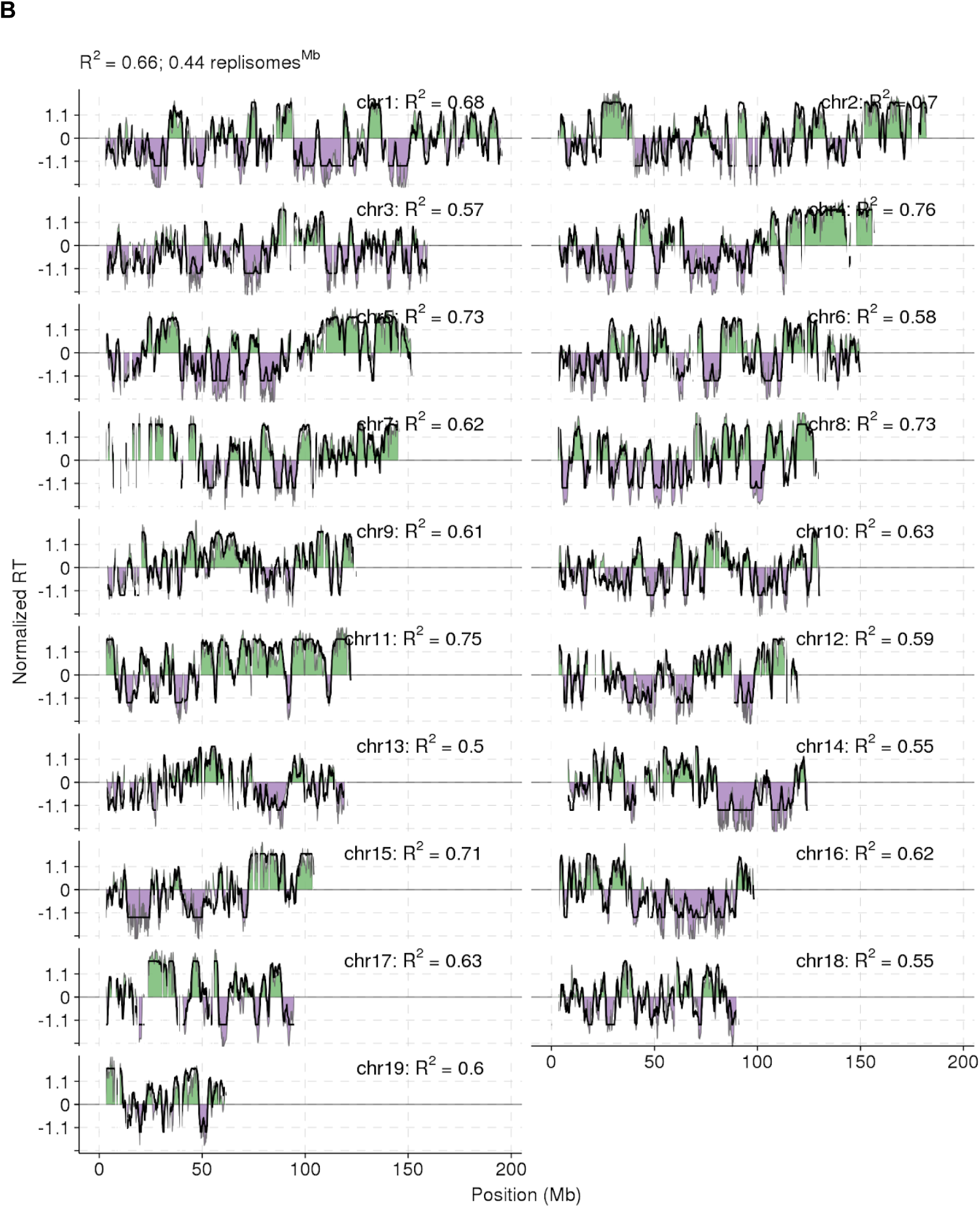
Modelling recapitulates experimental RT genome-wide. Experimental RT is shown as filled green (early replicating) and violet (late replicating) regions. The best-fitting RT-Sim predicted RT is shown as a solid black line. Per-chromosome Pearson correlation coefficients are shown. Both **(A)** Meiotic and **(B)** E14 ESC RT can be accurately modeled *in silico*. Both experimental and predicted RT are normalized by mean and standard deviation (s.d.); (RTnorm = (RT-mean(RT)) / s.d.(RT)).

**Figure S13.**
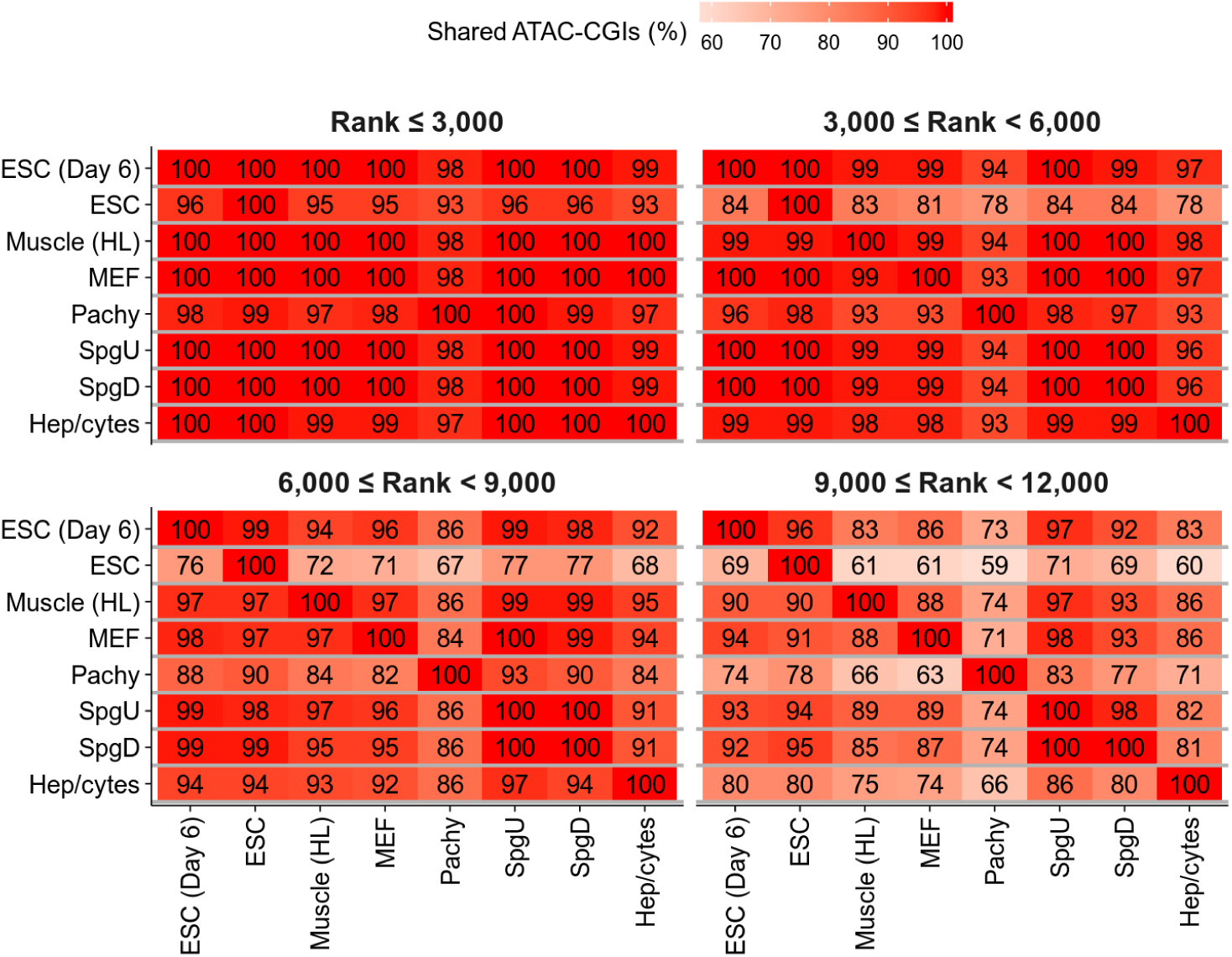
ATAC-Seq peaks at CpG islands are broadly conserved across diverse cell-types. We called peaks from 8 published ATAC-Seq experiments in a variety of mouse cell types. The strongest 12,000 peaks were split into 4 groups by strength. We counted the overlap of each peak subset with the top 12,000 ATAC-Seq peaks from each other cell-type. The percentage of peaks from each subset (y-axis) found in other cell types (x-axis) is shown. Datasets used are as follows: Hepatocytes (Hep/cytes; SRA accessions: SRR6813698,SRR6813699)(Li et al., 2019b), Differentiated cKIT+ Spermatogonia (SpgD; SRA accessions: SRR5956508)(Maezawa et al., 2018), Undifferentiated THY+ Spermatogonia (SpgU; SRA accessions: SRR5956504)(Maezawa et al., 2018), Pachytene Spermatocytes (Pachy; SRA accessions: SRR5956512)(Maezawa et al., 2018), Mouse Embryonic Fibroblasts (MEF; SRA accessions: SRR7048429, SRR7048430), Hindlimb Muscle Cells E14.5 (Muscle (HL); SRA accessions: SRR8104383,SRR8104391)(Castro et al., 2019), Embryonic Stem Cells (ESC; SRA accessions: SRR7048437,SRR7048438), Embryonic Stem Cells - 6 Days (ESC (Day 6); SRA accessions: SRR7048433,SRR7048434).

**Figure S14.**
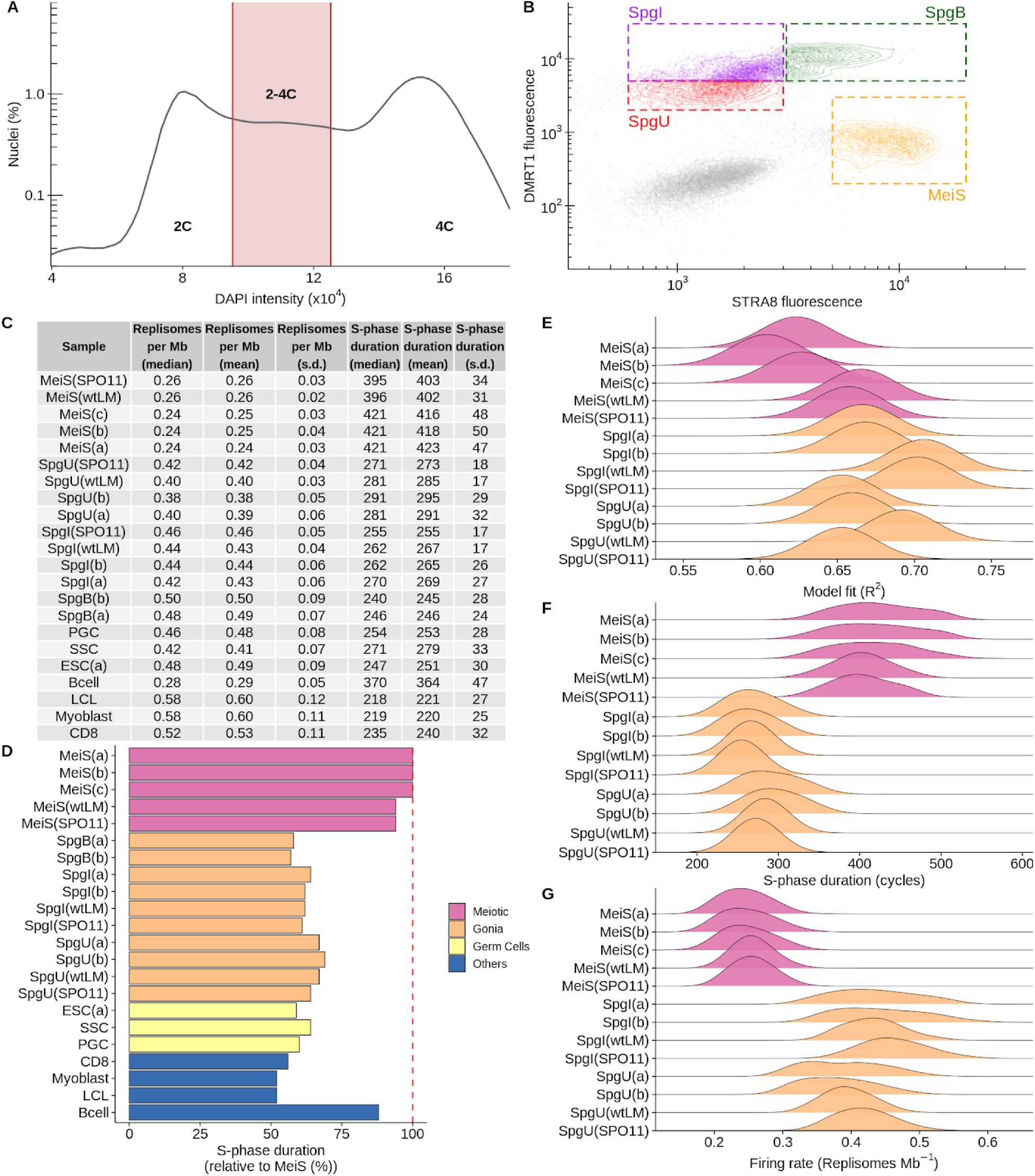
Meiotic S-phase duration is unaffected in *Spo11^-/-^* male mice. Meiotic S-phase nuclei, B-type spermatogonia and undifferentiated spermatogonia were isolated from a *Spo11^-/-^* male mouse testis (SPO11 in sample names) and a *wild-type* littermate (wtLM in sample names). Details for each sample are in Tables S1-3. (**A**) 2-4C nuclei were sorted and (**B**) gated by DMRT1 and STRA8 immunofluorescence signal. Gates were defined while sorting nuclei from the *Spo11^-/-^* mouse and retained, unchanged for sorting nuclei from the littermate. (**C**) Summary statistics for best-fitting RT-Sim models for all samples. (**D**) Comparison of S-phase duration across cell types. The median S-phase duration of good models for each cell-type is compared to meiotic S-phase. (**E**) The “fit” of RT-Sim models for *Spo11^-/-^* samples is comparable to the fit in other cell types. (**F**) The duration of meiotic S-phase is unaffected in *Spo11^-/-^* mice relative to the *wild-type* littermate control. Notably, both mice exhibit slightly shorter S-phase duration compared to the three *wild-type* replicates. This is likely because for the *Spo11^-/-^* and *wild-type* littermate, the sorting gates were defined in the *Spo11^-/-^* mouse and left unchanged when sorting nuclei from the littermate. Cellularity differences from the loss of *Spo11* may result in the capture of a slightly different population than when sorting gates are defined using *wild-type*. (**G**) The firing rate is similarly unaffected.

**Figure S15.**
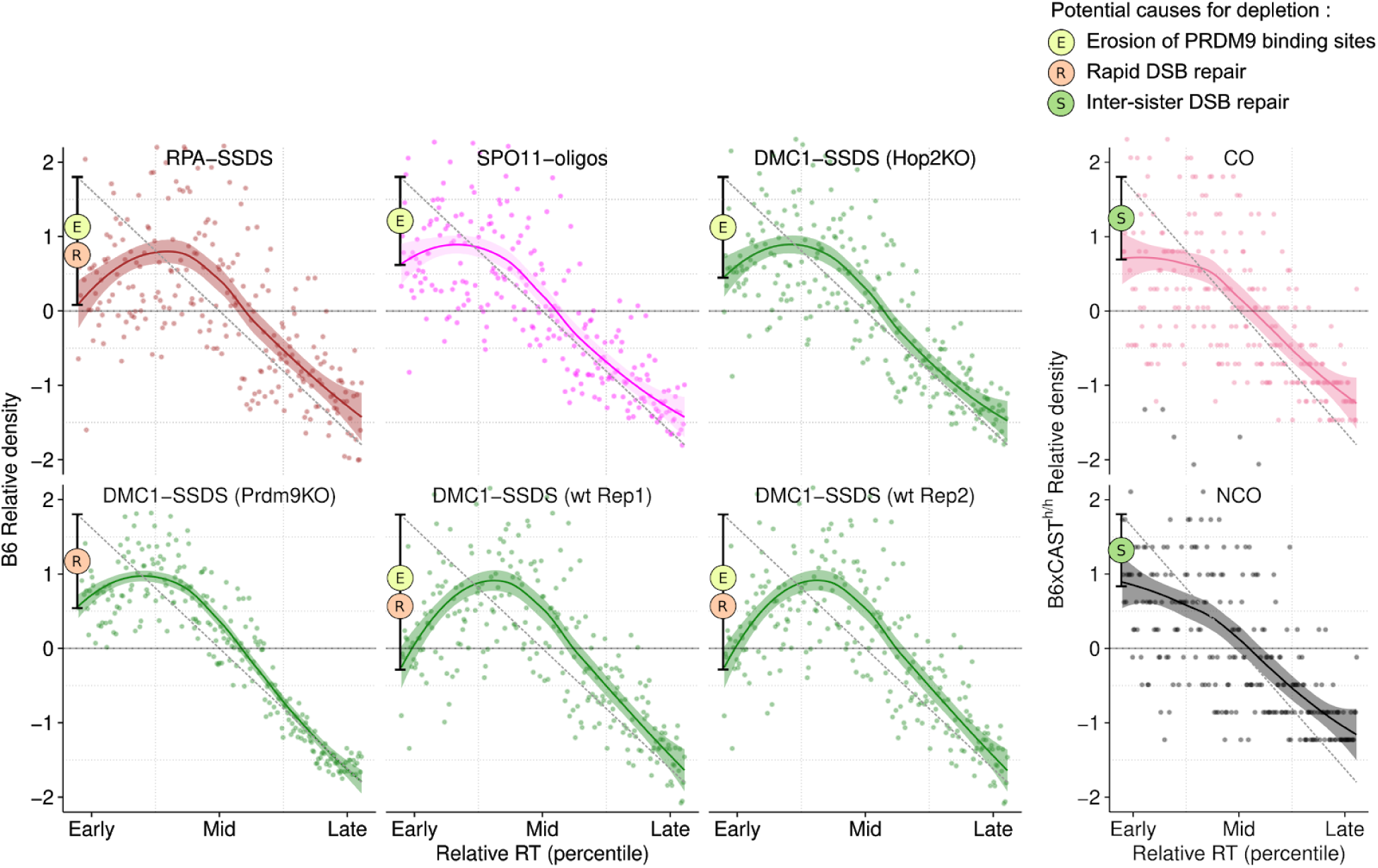
Rapid repair of DSBs in early replicating regions. SSDS using DMC1 exhibits a dip in the earliest replicating DNA. This is mirrored by SSDS using another ssDNA-binding protein, RPA. Thus, this dip does not simply reflect a property of DMC1 binding (two replicates shown). HOP2 is required for strand invasion and DSB repair. In *Hop2^-/-^* mice, DSBs are not repaired and therefore any effects resulting from repair dynamics should be absent. We find that the dip in the DMC1-SSDS signal in the earliest replicating DNA is abolished in *Hop2^-/-^* mice and that the DMC1-SSDS curve now only reflects the rate of DSB formation (like SPO11-oligo mapping; pink). Importantly, in mice lacking functional PRDM9, the DMC1-SSDS signal also dips at the earliest replicating DNA. PRDM9 binding site erosion does not affect PRDM9-independent DSB hotspots, therefore the dip is not as deep as in *wild-type* mice, where both erosion and rapid repair play a role. Both crossovers (COs) and Non-crossovers (NCO) are depleted in the earliest replicating DNA. For all figures, dots represent the average signal from all autosomal bins for each RT quantile (N = 250). Simulated RT from the T1 Meiocyte population (B6; Figure 3D) or simulated RT from a B6xCAST F1 hybrid (B6xCAST) is used for RT estimates. The solid lines depict the LOESS smoothed signal ± standard error (shaded). The dashed grey line is a projected linear correlation and the deviation from a linear correlation was manually added as a black range bar. The phenomena contributing to each dip are indicated by the colored circles (see key at top).

**Figure S16.**
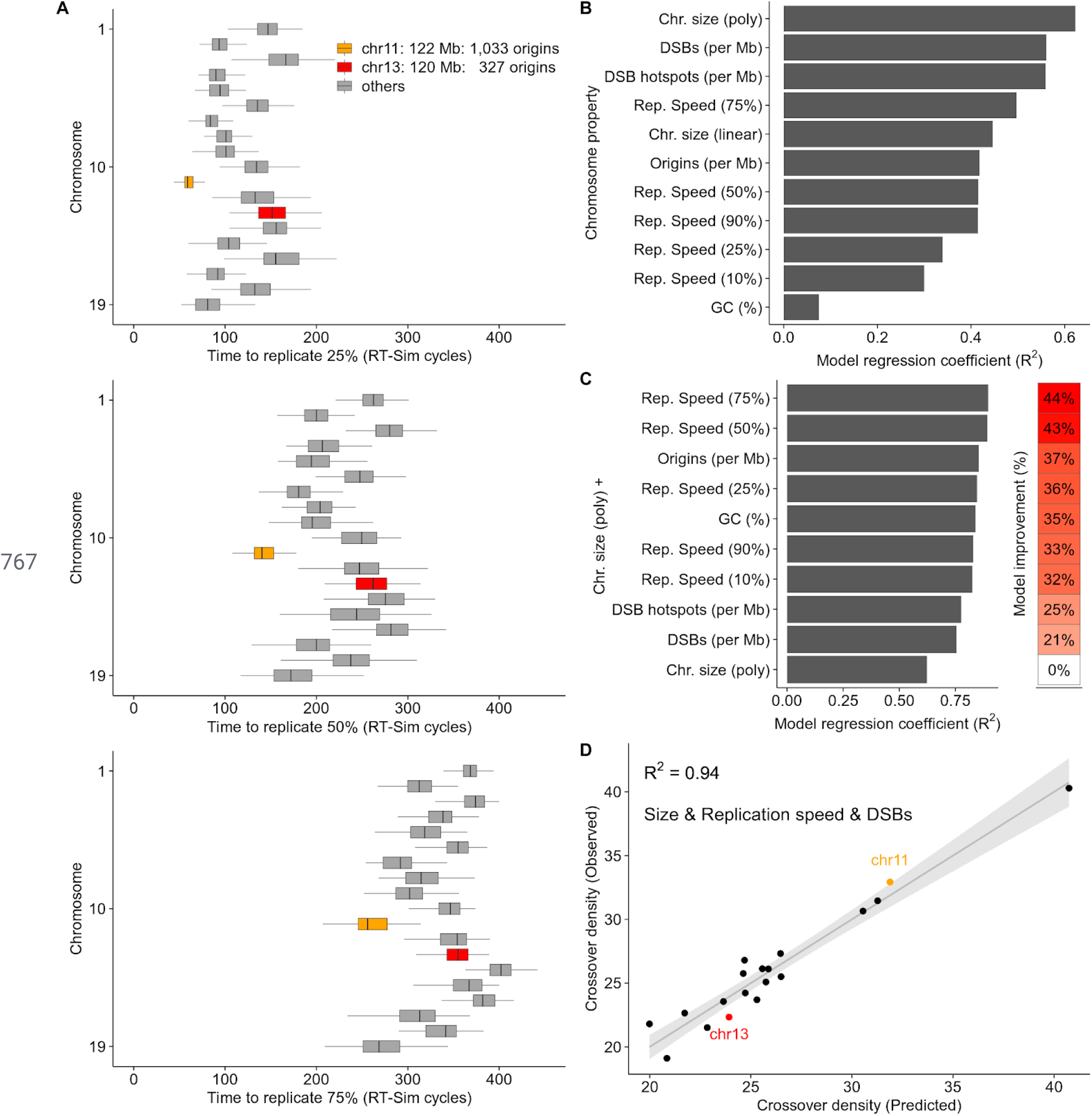
Replication speed differs among chromosomes and can predict crossover density. **(A)** Some chromosomes replicate faster than others; for example, chromosome 11 and 13 are a similar size but chromosome 11 replicates far earlier. Notably, chromosome 11 has over 3-times more replication origins. The time required to replicate 25% (top), 50% (middle) or 75% (bottom) of each chromosome is inferred from the best-fitting RT-Sim model. Boxplots indicate the range of times from 100 individual modelled cells. The median value for each chromosome is used as a measure of replication speed in subsequent panels. (**B**) We built simple regression models to predict crossover density from each individual property of chromosomes. DSBs are derived from DMC1-SSDS experiments in B6xCAST F1 mice (Smagulova et al., 2016). Crossovers in B6xCAST are taken from (Yin et al., 2019) and crossover density is calculated as total crossovers per Mb. Per-chromosome crossover density is best predicted by chromosome size (polynomial fit). This mirrors findings in other organisms (Murakami et al., 2020). (**C**) Next, we built a series of linear models of crossover density using chromosome size and each of the other chromosome properties. Replication speed and chromosome size together are a strong predictor of per-chromosome crossover density. The best combination (size + Rep.speed (75%)) yielded a 44% improvement on a model that used size alone and explained 90% of the variance in per-chromosome crossover density. Origin density and GC-content also improved the size-only models, but not as much as replication speed. (**D**) 94% of the variance in per-chromosome crossover density can be explained by adding DSB density to the best model from (C).

**Figure S17.**
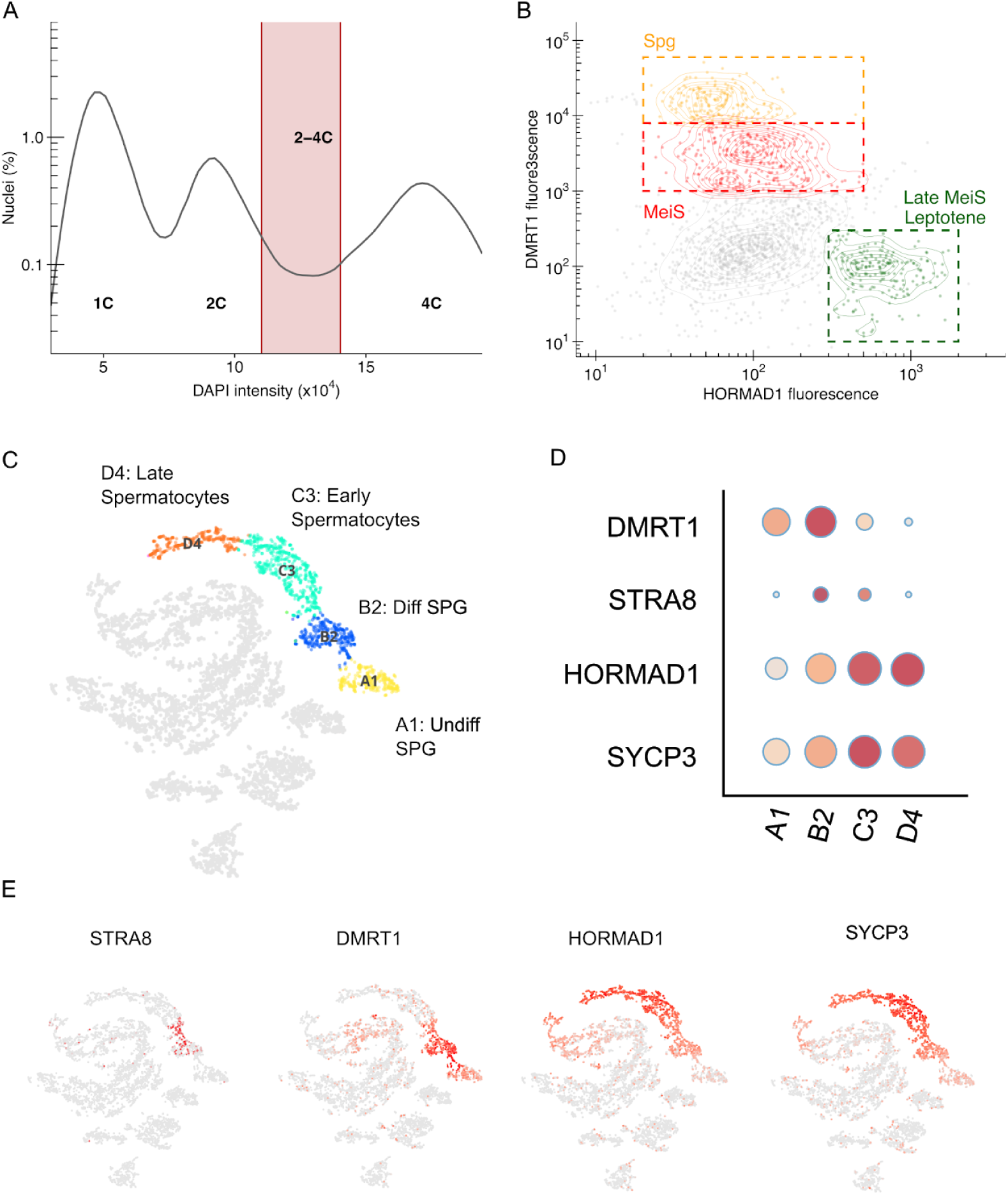
Isolation of human meiotic S-phase nuclei (A-B) Human meiotic S-phase and spermatogonia nuclei are isolated using fluorescence-activated nuclei sorting (Lam et al., 2019). (**A**) Replicating nuclei were isolated by gating 2-4C nuclei by DAPI content. (**B**) Putative meiotic nuclei were isolated using a combination of DMRT1 and HORMAD1. Antibodies to the STRA8 protein (used for sorting mouse meiotic S-phase nuclei) did not work for human. Unlike in mouse, the DMRT1 protein is expressed in meiotic S-phase (**C-E**). Thus, some spermatogonia may be present in the MeiS population. (**C**) *t*-Distributed Stochastic Neighbor Embedding (t-SNE) projection of the single-cell transcription atlas of human testes (Guo et al., 2018). Each dot represents a single cell. Colors represent defined clusters. (**D**) Quantification of genes-of-interest in four cell clusters. Circles represent the relative abundance of transcripts. Circle size represents the fraction of cells showing expression. Depth of color represents average gene expression. (**E**) Single-cell expression patterns of genes-of-interest. Depth of red color indicates expression level (Red = high; Grey = low).

**Figure S18.**
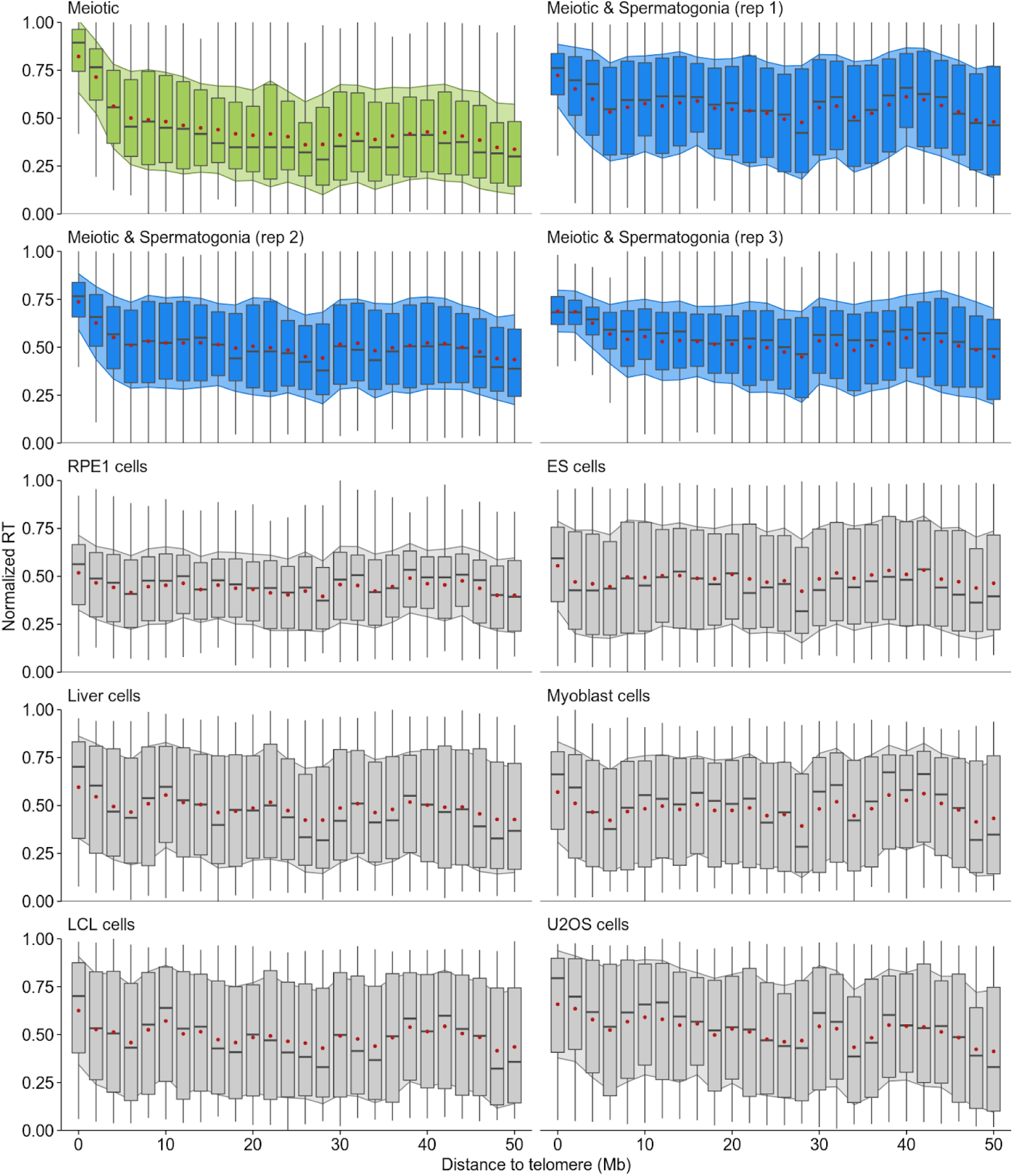
Early sub-telomeric replication is distinct to male meiosis. Consistently early replication in subtelomeric regions is only seen in the germline (green or blue) and not in any of the other cell types studied (grey). The pure meiotic S-phase population (green) exhibits the most pronounced signal. In the three samples which likely contain some spermatogonia (blue), early replication in subtelomeric DNA is seen, but less pronounced. This could imply that sub-telomeric DNA does not replicate notably early in spermatogonia. Boxplots depict the range of replication timing values in 2 Mb regions across the genome; the median is shown as a grey bar and the box designates the interquartile range. The mean (red dot) ± 1 standard deviation (filled shadow) is also shown for each interval. Details for each sample are in Table S1, S3.

## MMethods

### Tissue sources

C57BL/6J (stock 000664) and CAST/Eij (000928) mice were obtained from The Jackson Laboratory (Maine). Human testicular samples were obtained from a commercial source (Folio, Ohio) that provides testicular biopsies from normal tumor adjacent tissue or from deidentified donor testes from deceased individuals (obtained with the assistance of the Washington Regional Transplant Community).

### Mapping origins of replication using short nascent-strand capture followed by single-stranded DNA sequencing (Ori-SSDS)

For the short nascent strands (SNS) preparation we adapted the protocol from (Picard et al., 2014) to work with mouse tissue. Mice were euthanized and testes were retrieved. The tunica albuginea was removed and both testes were suspended in 10 ml of DNAzol (Thermo-Fisher catalog number 10503027) and transferred to a Dounce homogenizer. Tissue and cells were disrupted with 5-7 strokes of a loose-fitting pestle. The DNA was precipitated with 5 ml of EtOH and then spooled with a pipette tip and rinsed gently twice with EtOH 75% by sequential transfer into 15 ml tubes. DNA was air-dried for 5 min and then resuspended in 500 μl of 1X TEN buffer (Tris 10mM pH=8; EDTA 1mM; NaCl 100mM) + 80 U of RNAseOUT (Thermo-Fisher catalog number 10777019) (40 U/μl) at 4°C for at least 24h.

DNA was denatured at 95°C for 5 min, chilled on ice for 5 min and size-fractionated on 5 - 30% (w/v) linear sucrose gradients in TEN buffer in a Beckman Ultracentrifuge SW45 rotor for 18 h at 26,000 g at 20°C. 500 ul fractions were collected from the top of the gradient, precipitated by adding 1 ml of EtOH 100 % and 50 μl of NaOAc 3M pH 5.2 and resuspended in 25 μl Tris Buffer pH=8. Fractions were analyzed in a denaturing (50 mM NaOH, 1 mM EDTA) 1.2% agarose gel and the ones containing ssDNA ranging in size from 800 to 2000 nt were pooled. The sample was then heat-denatured for 5 min at 95°C and chilled on ice for 5 min and DNA was phosphorylated by Polynucleotide Kinase in 150 ul reactions in PNK Buffer (100 U of PNK ; 1X PNK buffer ; 1 mM ATP; 40U RNAseOUT). The reactions were incubated for 30 min at 37°C and heat-inactivated at 75°C for 15 min. DNA was extracted with Phenol/Chloroform, EtOH precipitated and resuspended in 50 μl Tris Buffer pH=8. Samples were heat-denatured again for 5 min and digested with lambda-exonuclease in 100 μl of custom lambda buffer (67 mM glycine-KOH pH= 8.8, 2.5 mM MgCl_2_, 50 μg/ml BSA) with 2.5 U to 10 U lambda-exonuclease/μg DNA (Thermo Scientific). 80U RNAseOUT, 5U of PNK and 1 mM ATP were added to the sample and incubated overnight at 37°C. Reactions were inactivated for 10 min at 75°C and DNA was purified with MinElute columns (QIAGEN), as indicated by the instruction manual. RNA primers were hydrolyzed by incubating the eluate for 30 min at 37°C in 0.25 N NaOH, followed by neutralization with acetic acid and purified again with MinElute columns. The sample was then sonicated in a Bioruptor UCD200 for 5 min 30 sec ON 30 sec OFF, “high” setting, and the library was prepared for SSDS as described (Brick et al., 2018b).

For one sample (wt_Rep2), separate libraries were made from three fractions from the sucrose gradient (Fraction 10 : 800-1,000 nt, Fraction 11 : 1,000-1,400 nt and Fraction 12 : 1,400-2,000 nt). Sequencing data were subsequently pooled.

To validate that the Ori-SSDS enrichment stems from RNA primed leading strands, we performed experiments where we hydrolyzed the RNA prior to lambda exonuclease treatment. This also necessitated changes to the above Ori-SSDS protocol as follows. Selected fractions from the sucrose gradient were split in two tubes. In one tube, leading strand RNA primers were hydrolyzed by incubating for 30 min at 37°C in 0.25 N NaOH, followed by neutralization with acetic acid. The other tube was incubated for 30 min at 37°C but in Tris Buffer pH=8. After neutralization, both tubes were heat denatured and spun through a Chromaspin TE-1000 size exclusion column (Clontech) to remove any small degradation products that could interfere with later steps. The eluate was then subjected to PNK treatment and subsequent steps outlined above. Two replicates were performed in parallel, with and without RNA hydrolysis (wt_Rep4, wt_Rep5).

Ori-SSDS in sperm was used as a negative control as there is no DNA replication. Sperm was retrieved from cauda and caput epididymides from 3 adult mice. The mix of somatic cells and sperm was resuspended in 0.1% SDS, 0.5% Triton X to selectively lyse somatic cells. Sperm was recovered by centrifugation and lysed in 6 M guanidinium, 30 mM sodium citrate (pH 7.0), 0.5% Sarkosyl, 0.20 mg/ml proteinase K and 0.3 M Beta-mercaptoethanol, and incubated at 55°C for 1h (Hossain et al., 1997). The DNA was precipitated with 2 volumes of isopropyl alcohol and then spooled with a pipette tip and rinsed gently twice with EtOH 75% by sequential transfer into 15 ml tubes. DNA was air-dried for 5 min and then resuspended in 500 μl of 1X TEN buffer (Tris 10mM pH=8; EDTA 1mM; NaCl 100mM) + 80 U of RNAseOUT (Thermo-Fisher catalog number 10777019) (40 U/μl) at 4°C for at least 24h.

### Identification of origins of replication

Sequencing reads from Ori-SSDS experiments were aligned to the reference genome (mouse = mm10; human = hg38) using bwa 0.7.12 and the single-stranded DNA pipeline (Brick et al., 2018b; Khil et al., 2012). Only fragments derived unambiguously from ssDNA (ssDNA_type1) were used in subsequent analyses. ssDNA fragments where either read1 or read2 had mapping quality <30 were discarded. Fragments from the mitochondrial chromosome and fragments in blacklisted regions of the genome were also discarded. Duplicate fragments were subsequently discarded.

We used the MACS algorithm for peak calling. To assure that data derived from Ori-SSDS fit the MACS peak model, we performed two operations: 1. The strand orientation of fragments was reversed prior to peak calling. 2. The data was passed to MACS as ssDNA fragment BED files (from SSDS pipeline). Peak calling was performed for each Ori-SSDS sample using MACS 2.1.2 and the following parameters : -g (hs *or* mm) -q 0.001 -extsize 2000 --nomodel --nolambda.

The final origin set for mouse was defined by merging origin calls from wt_Rep1,2,3. Origin intervals overlapping by ≥ 500 bp were merged. Origin efficiency was calculated in each sample using a method analogous to that for DMC1-SSDS hotspot mapping (Brick et al., 2018b). Briefly, each origin is re-centred to the midpoint of the Watson and Crick fragment distributions. Signal is estimated as the Crick-strand signal to the left of the origin center + the Watson-strand signal to the right of the origin center. The background is estimated by counting the remaining fragments at the left and right edges of MACS-defined origins (15%); Watson-strand fragments to the left of the origin center, Crick-strand fragments to the right. This background is then extrapolated to the entire origin and subtracted from the signal. Following this process, origins found in just one sample and origins that did not display the expected Crick/Watson asymmetry (Figure 1A) in any sample were discarded.

### Whole genome sequencing for replication timing

Nuclei from mouse or human testes were prepared as described in (Lam et al., 2019). To purify the populations of interest, a combination of intranuclear markers were used (Table S1, Figure 2).

Mouse ESC were obtained from Jackson labs: 129X1/SvJ-PRX-129X1 #1 mES cells derived from day 3.5 blastocysts of strain 129X1/SvJ (Stock Number 000691). The cells were cultured as recommended by Jackson labs. Exponentially growing cells were cross-linked by adding fresh 1% paraformaldehyde solution to the ES media for 10 minutes at room temperature. The cells were washed twice with 1× phosphate-buffered saline and scraped off the plates, pelleted and nuclei were prepared as described in (Lam et al., 2019). Replicating cells were sorted based DNA content.

Pure nuclei populations were resuspended in 300 ul of lysis buffer (SDS 1%, 10mM EDTA, 50mM, Tris pH=8) and then sonicated in a Bioruptor UCD200 for 20 min 30 sec ON 30 sec OFF, “high” setting. NaCl was added to a final concentration of 0.2M and the sample incubated overnight at 65°C to reverse DNA-protein crosslinks. 5U of Proteinase K (NEB Cat. No. P8107S) was added and the sample further incubated for 1h at 45°C. DNA was purified with MinElute columns (QIAGEN), as indicated by the instruction manual. Whole genome sequencing libraries were prepared from 100 ng of purified DNA using the KAPA Hyper Prep Kit (Cat. No. KK8502) following the manufacturer’s instructions.

### Inferring replication timing from whole genome sequencing (WGS) data

The mem algorithm of bwa 0.7.12 (Li, 2013; Li and Durbin, 2009) was used to align each WGS dataset to the reference genome (mm10 for mouse; hg38 for human). WGS from a non-replicating population was used to calibrate the expected GC-biases from different sequencing instruments (Illumina HiSeq 2000, Illumina HiSeq 2500, Illumina HiSeq X, Illumina NextSeq 500). Briefly, these calibration files were generated by examining a subset of perfectly mappable genomic loci. Each locus is a 101 bp region with the following properties:

1. Full mappability of all bases within the 101 bp window. To determine mappability, we generated a fastq file of pseudoreads of the requisite length from the reference genome fasta file. One pseudo-read was generated per bp; all bases were assigned a q-score of 60. Pseudo-reads were mapped to the reference genome using bwa mem 0.7.12 and the coverage at each base in the genome was assessed.
2. The window does not overlap with sequencing gaps (UCSC Gaps track), segmental duplications (UCSC SegDups track), high copy repeats (UCSC repeatmasker track) or otherwise blacklisted genomic regions (Amemiya et al., 2019; Kent et al., 2002).

The samples used to generate GC-correction files were the 2C_MMC_ and 2C_HSC_ samples (Table S1) for Hi-Seq X and Hi-Seq 2500, respectively. Samples used to generate GC-correction files for published data are detailed in Table S2 below.

The GC content (rounded to 1%) and WGS sequencing coverage of each 101 bp window was assessed. The median coverage for all windows of a given GC content was used as a correction factor. These correction factors vary substantially across the different sequencing platforms used. 2% of all qualifying 101 bp autosomal windows were used to calculate the GC correction coefficients.

For each RT-Seq experiment, the WGS coverage in 101-bp windows in the genome was calculated and corrected for GC biases using the correction coefficient for the appropriate sequencing platform. Replication timing was calculated as the log_2_ ratio of corrected sequencing coverage at a locus to the genome-wide median (method was adapted from (Koren et al., 2014)). RT-Seq coverage was smoothed in 500 Kb windows with a 50 Kb step.

### Inferring replication timing from published data

Whole genome sequencing data were obtained from the sources listed in Table S2. Sequencing data were aligned and replication timing profiles were inferred as described above.

### Modelling DNA replication

DNA replication was modelled *in-silico* using custom R code (Brick, 2020). Replication was simulated in a haploid genome. Non-autosomal chromosomes were excluded. The initial wave of origin firing was assumed to be synchronous. *k* replication origins were selected at random and without replacement, using the origin efficiency as a selection weight. At each origin, a left-ward and a right-ward replication fork was established, and replication was then simulated bi-directionally outward from each origin in 10 Kb steps. After each step, we assessed which replication forks have collided with another fork and which forks reached the end of the chromosome. These forks were terminated. For every two terminated forks, we allowed replication to begin from a new, randomly chosen origin (as above, weighted by origin efficiency). Origins that have previously fired and origins in regions that have already been replicated were not considered for selection. This iterative process continued until the genome was fully replicated or for a defined number of cycles. To simulate replication in a population, this simulation was performed in *n* haploid genomes and the results were combined.

### Hyperparameter search for replication models

To assess the best-fitting parameters of DNA replication *in-silico*, we performed a grid-hyperparameter search. A hyperparameter search was performed for each pair-wise combination of RT and replication origins. We optimized for a scoring function defined as the Pearson correlation coefficient between the experimentally-determined replication timing and the *in-silico* simulated replication timing. A population of 250 replicating haploid genomes was used for the hyperparameter search. The grid-search generated an RT-Sim model for each combination of the following parameters:

a. Origin density; R_d_ = 0.02 to 0.4 in 0.01 steps; 0.45 to 0.6 in 0.05 steps
b. Modelling runtime; N_epochs_ = 10 to 7500 in steps of 10
c. Replicating cells in population; P (%) = 10,30,50,70,90
d. Use origin efficiency; E = TRUE / FALSE (if FALSE, all origins were equally weighted for “random” selection)
e. fine-grained search yielded multiple models with similar scores, therefore in assessing results, the top *g*% of models were examined.

### Generating simulated replication timing

The hyperparameters of the top-scoring RT-Sim model were used to generate an RT-Sim model from 500 haploid genomes and run it to completion. Simulated replication timing was then defined as the mean replication time across all simulations for each 10 Kb window in the genome.

### Sources and processing of annotation data for the mouse genome

CpG islands in the mouse genome were obtained from the Irizarry lab at Harvard University, USA (http://www.haowulab.org/software/makeCGI/model-based-cpg-islands-mm9.txt) (Wu et al., 2010). UCSC liftover was used to convert mm9 to mm10 genome coordinates.

Meiotic DSB hotspots in C57BL6J mice were obtained from columns 1,2,3,6 of Supplementary File 41586_2018_492_MOESM3_ESM.zip (Brick et al., 2018a). H3K4m3 ChIP-Seq signals at DSB hotspots in C57BL6J mice (Baker et al., 2014) was obtained from columns 1,2,3,50 of Supplementary File 41586_2018_492_MOESM3_ESM.zip (Brick et al., 2018a). Genome-wide Spo11-associated oligo mapping data in C57BL6 mice (Lange et al., 2016) were obtained in processed form from columns 1,2,3,55 of Supplementary File 41586_2018_492_MOESM3_ESM.zip (Brick et al., 2018a). DSB hotspots in *Prdm9^-^*^/-^ mice (Brick et al., 2012) were obtained from GSM2664291 (Brick et al., 2018a). DSB hotspots from mice homozygous for a humanized PRDM9 zinc-finger array (B6^h/h^) were obtained from the GEO (accession: GSM2049306; file: GSM2049306_dmc1hotspots_B6.PRDM9hh.txt.gz) (Davies et al., 2016). B6^h/h^ hotspots overlapping hotspots from *Prdm9*^-/-^ were discarded. H3K4m3 ChIP-Seq peaks in B6^h/h^ mice were obtained from the GEO (accession: GSM1904284; file GSM1904284_H3K4me3.ForceCalledValues.B6.PRDM9hh.txt) (Davies et al., 2016). Crossovers and non-crossovers defined by the humanized variant of mouse PRDM9 were obtained from supplementary file 3 (41467_2019_11675_MOESM6_ESM.csv) and 4 (41467_2019_11675_MOESM7_ESM.csv), respectively, from (Li et al., 2019a). Affinity-Seq data describing the *in-vitro* binding preferences of the C57BL6J PRDM9 variant were obtained from the Sequence Read Archive (accession: SRR1976005) (Walker et al., 2015). Control data used for Affinity-Seq peak calling was obtained from the Sequence Read Archive (accession: SRR1976003 & SRR1976004) (Walker et al. 2015). Affinity-Seq and control reads were aligned to the reference mm10 genome using bwa mem 0.7.12. Affinity-Seq peak calling was performed using MACS 2.1.0.20150731 with the following arguments: -g mm -bw 1000 --keep-dup all --slocal 5000 -q 0.05. Affinity-Seq peak strength was determined as Affinity-Seq coverage less Control DNA coverage at Affinity-Seq peaks. Hi-C data from mouse zygotene cells was downloaded from the GEO (GEO accession: GSE122622; file: GSE122622_zygotene_overall.hic) (Patel et al., 2019). The Juicer tools 1.5.6 eigenvector program (Durand et al., 2016) was used to extract the first eigenvector from the .hic file, using the following arguments: KR, BP, binsize = 100000, -p. H3K9m2 (accession: SRR1975998) and H3K9m3 (accession: SRR1585300) ChIP-Seq data (Walker et al., 2015) from spermatocytes were obtained from the SRA. Reads were aligned to the reference mm10 genome using bwa mem 0.7.12.

ATAC-Seq data were obtained from the published datasets described in Supplementary Figure 9. Each dataset was aligned to the mouse mm10 reference genome using bwa mem 0.7.12. Peak calling was performed for each dataset using MACS 2.1.2 without a control and using default arguments. ATAC-Seq peaks coinciding with CpG-islands were defined for use on Supplementary Figure 9.

### Mapping Okazaki-fragment sequencing data

Data from Okazaki-fragment sequencing in mouse ES-cells were obtained from SRA accession SRR7535256 (Petryk et al., 2018). Sequencing reads were aligned to the mm10 reference genome using the mem algorithm of bwa 0.7.12 (Li, 2013; Li and Durbin, 2009).

### Obtaining published SNS-Seq replication origins data

Processed peak calls for MEFs and ESCs were obtained from GEO accession GSE99741 (Almeida et al., 2018); MEFs: GSM2651107_MEFs_WT_repl_I_peaks.bigBed; ESCs: GSM2651111_mES_WT_repl_I_peaks.bigBed. Processed peak calls for ESCs from (Cayrou et al., 2015) were obtained from GEO accession GSE68347; GSE68347_Initiation_Sites.bedGraph.gz. The UCSC liftover tool was used to convert mm9 to mm10 coordinates.

### Mapping DNA DSB hotspots from DMC1-SSDS experiments

A testicular biopsy was obtained from a commercial source (Folio, Ohio). Genotyping for PRDM9 alleles using Sanger sequencing (Pratto et al., 2014) revealed that this individual did not carry a copy of the most common PRDM9 variant in humans (PRDM9_A_). Instead, this individual was heterozygous for the PRDM9_C_ and PRDM9_L4_ alleles (Berg et al., 2010). A DMC1-SSDS library was prepared using the protocol in (Brick et al., 2018b; Pratto et al., 2014). Published DMC1-SSDS data were also obtained for two human PRDM9_A_-homozygous males (GEO accessions: GSM1447325 & GSM1447328) (Pratto et al., 2014). Sequencing reads were aligned to the human hg38 reference genome using bwa 0.7.12 and the single-stranded DNA pipeline (Brick et al., 2018b; Khil et al., 2012). Hotspots were called using the DMC1-SSDS DSB hotspot calling pipeline (Brick et al., 2018b).

### Crossover and non-crossover data for humans

Crossover data for human males and females were obtained from the supplementary data files S1 and S2, respectively, from (Halldorsson et al., 2019). Non-crossover data were obtained from supplementary file S2 from (Halldorsson et al., 2016). NCOs from the ChIP-based approach and the sequencing-based approach were merged.

### Data analysis tools and versions

BEDtools v.2.29.2 (Quinlan and Hall, 2010) BWA 0.7.12 (Li, 2013; Li and Durbin, 2009).

DeepTools v.3.0.1 (Ramírez et al., 2014) Juicer v.1.5.6 (Durand et al., 2016)

MACS v.2.1.0.20150731 (Zhang et al., 2008)

MACS v.2.1.2 (Zhang et al., 2008)

Nextflow v.20.01.0 (Di Tommaso et al., 2017)

Picard v.2.9.2 (Broad Institute of MIT and Harvard, 2018) R v.3.6.0 (R Core Team, 2014)

SAMtools v.1.9 (Li et al., 2009)

SRA toolkit v.2.9.2 (Leinonen et al., 2011) UCSC toolkit v.385 (Kent et al., 2010)

## Notes

### Competing Interest Statement

The authors have declared no competing interest.

## References

Almeida, R., Fernández-Justel, J.M., Santa-María, C., Cadoret, J.-C., Cano-Aroca, L., Lombraña, R., Herranz, G., Agresti, A., and Gómez, M. (2018). Chromatin conformation regulates the coordination between DNA replication and transcription. Nat. Commun. 9, 1590.

Amemiya, H.M., Kundaje, A., and Boyle, A.P. (2019). The ENCODE Blacklist: Identification of Problematic Regions of the Genome. Sci. Rep. 9, 9354.

Baker, C.L., Walker, M., Kajita, S., Petkov, P.M., and Paigen, K. (2014). PRDM9 binding organizes hotspot nucleosomes and limits Holliday junction migration. Genome Res. 24, 724–732.

Baudat, F., Buard, J., Grey, C., Fledel-Alon, A., Ober, C., Przeworski, M., Coop, G., and de Massy, B. (2010). PRDM9 is a major determinant of meiotic recombination hotspots in humans and mice. Science 327, 836–840.

Bellve, A.R., Cavicchia, J.C., and Millette, C.F. (1977). Spermatogenic cells of the prepuberal mouse: isolation and morphological characterization. J. Cell Biol.

Bennett, M.D., Smith, J.B., and Riley, R. (1972). The effects of polyploidy on meiotic duration and pollen development in cereal anthers. Proceedings of the Royal Society of London. Series B. Biological Sciences 181, 81–107.

Berg, I.L., Neumann, R., Lam, K.-W.G., Sarbajna, S., Odenthal-Hesse, L., May, C.A., and Jeffreys, A.J. (2010). PRDM9 variation strongly influences recombination hot-spot activity and meiotic instability in humans. Nat. Genet. 42, 859–863.

Bergerat, A., de Massy, B., Gadelle, D., Varoutas, P.C., Nicolas, A., and Forterre, P. (1997). An atypical topoisomerase II from Archaea with implications for meiotic recombination. Nature 386, 414–417.

Bielinsky, A.K., and Gerbi, S.A. (1998). Discrete start sites for DNA synthesis in the yeast ARS1 origin. Science 279, 95–98.

Bishop, D.K., Park, D., Xu, L., and Kleckner, N. (1992). DMC1: a meiosis-specific yeast homolog of E. coli recA required for recombination, synaptonemal complex formation, and cell cycle progression. Cell 69, 439–456.

Blitzblau, H.G., Chan, C.S., Hochwagen, A., and Bell, S.P. (2012). Separation of DNA replication from the assembly of break-competent meiotic chromosomes. PLoS Genet. 8, e1002643.

Borde, V., Goldman, A.S., and Lichten, M. (2000). Direct coupling between meiotic DNA replication and recombination initiation. Science 290, 806–809.

Boulton, A., Myers, R.S., and Redfield, R.J. (1997). The hotspot conversion paradox and the evolution of meiotic recombination. Proc. Natl. Acad. Sci. U. S. A. 94, 8058–8063.

Brick, K. (2020). Simulating replication timing in the mouse and human genomes (Github).

Brick, K., Smagulova, F., Khil, P., Camerini-Otero, R.D., and Petukhova, G.V. (2012). Genetic recombination is directed away from functional genomic elements in mice. Nature 485, 642–645.

Brick, K., Thibault-Sennett, S., Smagulova, F., Lam, K.-W.G., Pu, Y., Pratto, F., Camerini-Otero, R.D., and Petukhova, G.V. (2018a). Extensive sex differences at the initiation of genetic recombination. Nature 561, 338–342.

Brick, K., Pratto, F., Sun, C.-Y., Camerini-Otero, R.D., and Petukhova, G. (2018b). Analysis of Meiotic Double-Strand Break Initiation in Mammals. In Methods in Enzymology, (Academic Press), pp. 391–418.

Broach, J.R., Li, Y.Y., Feldman, J., Jayaram, M., Abraham, J., Nasmyth, K.A., and Hicks, J.B. (1983). Localization and sequence analysis of yeast origins of DNA replication. Cold Spring Harb. Symp. Quant. Biol. 47 *Pt* *2*, 1165–1173.

Broad Institute of MIT and Harvard (2018). Picard tools (Broad Institute of MIT and Harvard).

Buard, J., Rivals, E., Dunoyer de Segonzac, D., Garres, C., Caminade, P., de Massy, B., and Boursot, P. (2014). Diversity of Prdm9 zinc finger array in wild mice unravels new facets of the evolutionary turnover of this coding minisatellite. PLoS One 9, e85021.

Callan, H.G. (1973). DNA Replication in the Chromosomes of Eukaryotes. In Molecular Cytogenetics, B.A. Hamkalo, and J. Papaconstantinou, eds. (Boston, MA: Springer New York), pp. 31–47.

Castro, J.P.L., Yancoskie, M.N., Marchini, M., Belohlavy, S., Hiramatsu, L., Kučka, M., Beluch, W.H., Naumann, R., Skuplik, I., Cobb, J., et al. (2019). An integrative genomic analysis of the Longshanks selection experiment for longer limbs in mice. eLife 8.

Cayrou, C., Grégoire, D., Coulombe, P., Danis, E., and Méchali, M. (2012). Genome-scale identification of active DNA replication origins. Methods 57, 158–164.

Cayrou, C., Ballester, B., Peiffer, I., Fenouil, R., Coulombe, P., Andrau, J.-C., van Helden, J., and Méchali, M. (2015). The chromatin environment shapes DNA replication origin organization and defines origin classes. Genome Res. 25, 1873–1885.

Cha, R.S., and Kleckner, N. (2002). ATR homolog Mec1 promotes fork progression, thus averting breaks in replication slow zones. Science 297, 602–606.

Cha, R.S., Weiner, B.M., Keeney, S., Dekker, J., and Kleckner, N. (2000). Progression of meiotic DNA replication is modulated by interchromosomal interaction proteins, negatively by Spo11p and positively by Rec8p. Genes Dev. 14, 493–503.

Chagin, V.O., Casas-Delucchi, C.S., Reinhart, M., Schermelleh, L., Markaki, Y., Maiser, A., Bolius, J.J., Bensimon, A., Fillies, M., Domaing, P., et al. (2016). 4D Visualization of replication foci in mammalian cells corresponding to individual replicons. Nat. Commun. 7, 11231.

Chandley, A.C. (1986). A model for effective pairing and recombination at meiosis based on early replicating sites (R-bands) along chromosomes. Hum. Genet. 72, 50–57.

Davies, B., Hatton, E., Altemose, N., Hussin, J.G., Pratto, F., Zhang, G., Hinch, A.G., Moralli, D., Biggs, D., Diaz, R., et al. (2016). Re-engineering the zinc fingers of PRDM9 reverses hybrid sterility in mice. Nature 530, 171–176.

Diagouraga, B., Clément, J.A.J., Duret, L., Kadlec, J., de Massy, B., and Baudat, F. (2018). PRDM9 Methyltransferase Activity Is Essential for Meiotic DNA Double-Strand Break Formation at Its Binding Sites. Mol. Cell.

Dileep, V., and Gilbert, D.M. (2018). Single-cell replication profiling to measure stochastic variation in mammalian replication timing. Nat. Commun. 9, 427.

Di Tommaso, P., Chatzou, M., Floden, E.W., Barja, P.P., Palumbo, E., and Notredame, C. (2017). Nextflow enables reproducible computational workflows. Nat. Biotechnol. 35, 316–319.

Durand, N.C., Shamim, M.S., Machol, I., Rao, S.S.P., Huntley, M.H., Lander, E.S., and Aiden, E.L. (2016). Juicer Provides a One-Click System for Analyzing Loop-Resolution Hi-C Experiments. Cell Syst 3, 95–98.

Duret, L., and Galtier, N. (2009). Biased gene conversion and the evolution of mammalian genomic landscapes. Annu. Rev. Genomics Hum. Genet. 10, 285–311.

Foulk, M.S., Urban, J.M., Casella, C., and Gerbi, S.A. (2015). Characterizing and controlling intrinsic biases of lambda exonuclease in nascent strand sequencing reveals phasing between nucleosomes and G-quadruplex motifs around a subset of human replication origins. Genome Res. 25, 725–735.

Fu, H., Besnard, E., Desprat, R., Ryan, M., Kahli, M., Lemaitre, J.-M., and Aladjem, M.I. (2014). Mapping replication origin sequences in eukaryotic chromosomes. Curr. Protoc. Cell Biol. 65, 22–20.

Ghosal, S.K., and Mukherjee, B.B. (1971). THE CHRONOLOGY OF DNA SYNTHESIS, MEIOSIS AND SPERMIOGENESIS IN THE MALE MOUSE AND GOLDEN HAMSTER. Can. J. Genet. Cytol. 13, 672–682.

Gindin, Y., Valenzuela, M.S., Aladjem, M.I., Meltzer, P.S., and Bilke, S. (2014). A chromatin structure-based model accurately predicts DNA replication timing in human cells. Mol. Syst. Biol. 10, 722.

Goodsell, D.S., and Dickerson, R.E. (1994). Bending and curvature calculations in B-DNA. Nucleic Acids Res. 22, 5497–5503.

Grant, G.D., Kedziora, K.M., Limas, J.C., Cook, J.G., and Purvis, J.E. (2018). Accurate delineation of cell cycle phase transitions in living cells with PIP-FUCCI. Cell Cycle 17, 2496–2516.

Guo, J., Grow, E.J., Mlcochova, H., Maher, G.J., Lindskog, C., Nie, X., Guo, Y., Takei, Y., Yun, J., Cai, L., et al. (2018). The adult human testis transcriptional cell atlas. Cell Res. 28, 1141–1157.

Halldorsson, B.V., Hardarson, M.T., Kehr, B., Styrkarsdottir, U., Gylfason, A., Thorleifsson, G., Zink, F., Jonasdottir, A., Jonasdottir, A., Sulem, P., et al. (2016). The rate of meiotic gene conversion varies by sex and age. Nat. Genet. 48, 1377–1384.

Halldorsson, B.V., Palsson, G., Stefansson, O.A., Jonsson, H., Hardarson, M.T., Eggertsson, H.P., Gunnarsson, B., Oddsson, A., Halldorsson, G.H., Zink, F., et al. (2019). Characterizing mutagenic effects of recombination through a sequence-level genetic map. Science 363.

Handel, M.A., Eppig, J.J., and Schimenti, J.C. (2014). Applying “Gold Standards” to In-Vitro-Derived Germ Cells. Cell 157, 1257–1261.

Higgins, J.D., Perry, R.M., Barakate, A., Ramsay, L., Waugh, R., Halpin, C., Armstrong, S.J., and Franklin, F.C.H. (2012). Spatiotemporal asymmetry of the meiotic program underlies the predominantly distal distribution of meiotic crossovers in barley. Plant Cell 24, 4096–4109.

Hinch, A.G., Zhang, G., Becker, P.W., Moralli, D., Hinch, R., Davies, B., Bowden, R., and Donnelly, P. (2019). Factors influencing meiotic recombination revealed by whole-genome sequencing of single sperm. Science 363, eaau8861.

Hiratani, I., Ryba, T., Itoh, M., Rathjen, J., Kulik, M., Papp, B., Fussner, E., Bazett-Jones, D.P., Plath, K., Dalton, S., et al. (2010). Genome-wide dynamics of replication timing revealed by in vitro models of mouse embryogenesis. Genome Res. 20, 155–169.

Holm, P.B. (1977). The premeiotic DNA replication of euchromatin and heterochromatin in Lilium longiflorum (Thunb.). Carlsberg Res. Commun. 42, 249–281.

Hossain, A.M., Rizk, B., Behzadian, A., and Thorneycroft, I.H. (1997). Modified guanidinium thiocyanate method for human sperm DNA isolation. Mol. Hum. Reprod. 3, 953–956.

Huppert, J.L., and Balasubramanian, S. (2005). Prevalence of quadruplexes in the human genome. Nucleic Acids Res. 33, 2908–2916.

Jodkowska, K., Pancaldi, V., Almeida, R., Rigau, M., Graña-Castro, O., Fernández-Justel, J.M., Rodríguez-Acebes, S., Rubio-Camarillo, M., Pau, E.C.S., Pisano, D., et al. (2019). Three-dimensional connectivity and chromatin environment mediate the activation efficiency of mammalian DNA replication origins.

Joshi, N., Brown, M.S., Bishop, D.K., and Börner, G.V. (2015). Gradual implementation of the meiotic recombination program via checkpoint pathways controlled by global DSB levels. Mol. Cell 57, 797–811.

Kaback, D.B., Guacci, V., Barber, D., and Mahon, J.W. (1992). Chromosome size-dependent control of meiotic recombination. Science 256, 228–232.

Keeney, S., Giroux, C.N., and Kleckner, N. (1997). Meiosis-specific DNA double-strand breaks are catalyzed by Spo11, a member of a widely conserved protein family. Cell 88, 375–384.

Kelly, T., and Callegari, A.J. (2019). Dynamics of DNA replication in a eukaryotic cell. Proc. Natl. Acad. Sci. U. S. A. 116, 4973–4982.

Kent, W.J., Sugnet, C.W., Furey, T.S., Roskin, K.M., Pringle, T.H., Zahler, A.M., and Haussler, D. (2002). The human genome browser at UCSC. Genome Res. 12, 996–1006.

Kent, W.J., Zweig, A.S., Barber, G., Hinrichs, A.S., and Karolchik, D. (2010). BigWig and BigBed: enabling browsing of large distributed datasets. Bioinformatics 26, 2204–2207.

Khil, P.P., Smagulova, F., Brick, K.M., Camerini-Otero, R.D., and Petukhova, G.V. (2012). Sensitive mapping of recombination hotspots using sequencing-based detection of ssDNA. Genome Res. 22, gr – 130583.

Klein, K.N., Zhao, P.A., Lyu, X., Bartlett, D.A., Singh, A., Tasan, I., Watts, L.P., Hiraga, S.-I., Natsume, T., Zhou, X., et al. (2019). Replication timing maintains the global epigenetic state in human cells.

Kofman-Alfaro, S., and Chandley, A.C. (1970). Meiosis in the male mouse. An autoradiographic investigation. Chromosoma 31, 404–420.

Kojima, M.L., de Rooij, D.G., and Page, D.C. (2019). Amplification of a broad transcriptional program by a common factor triggers the meiotic cell cycle in mice. Elife 8.

Koren, A., Polak, P., Nemesh, J., Michaelson, J.J., Sebat, J., Sunyaev, S.R., and McCarroll, S.A. (2012). Differential relationship of DNA replication timing to different forms of human mutation and variation. Am. J. Hum. Genet. 91, 1033–1040.

Koren, A., Handsaker, R.E., Kamitaki, N., Karlić, R., Ghosh, S., Polak, P., Eggan, K., and McCarroll, S.A. (2014). Genetic variation in human DNA replication timing. Cell 159, 1015–1026.

Kulakovskiy, I.V., Vorontsov, I.E., Yevshin, I.S., Sharipov, R.N., Fedorova, A.D., Rumynskiy, E.I., Medvedeva, Y.A., Magana-Mora, A., Bajic, V.B., Papatsenko, D.A., et al. (2018). HOCOMOCO: towards a complete collection of transcription factor binding models for human and mouse via large-scale ChIP-Seq analysis. Nucleic Acids Res. 46, D252–D259.

Lam, K.-W.G., Brick, K., Cheng, G., Pratto, F., and Camerini-Otero, R.D. (2019). Cell-type-specific genomics reveals histone modification dynamics in mammalian meiosis. Nat. Commun. 10, 3821.

Lange, J., Yamada, S., Tischfield, S.E., Pan, J., Kim, S., Zhu, X., Socci, N.D., Jasin, M., and Keeney, S. (2016). The Landscape of Mouse Meiotic Double-Strand Break Formation, Processing, and Repair. Cell 167, 695–708.e16.

Lee, B., and Amon, A. (2001). Meiosis: how to create a specialized cell cycle. Curr. Opin. Cell Biol. 13, 770–777.

Leinonen, R., Sugawara, H., Shumway, M., and International Nucleotide Sequence Database Collaboration (2011). The sequence read archive. Nucleic Acids Res. 39, D19–D21.

Li, H. (2013). Aligning sequence reads, clone sequences and assembly contigs with BWA-MEM.

Li, H., and Durbin, R. (2009). Fast and accurate short read alignment with Burrows-Wheeler transform. Bioinformatics 25, 1754–1760.

Li, H., Handsaker, B., Wysoker, A., Fennell, T., Ruan, J., Homer, N., Marth, G., Abecasis, G., Durbin, R., and 1000 Genome Project Data Processing Subgroup (2009). The Sequence Alignment/Map format and SAMtools. Bioinformatics 25, 2078–2079.

Li, N., Lam, W.H., Zhai, Y., Cheng, J., Cheng, E., Zhao, Y., Gao, N., and Tye, B.-K. (2018). Structure of the origin recognition complex bound to DNA replication origin. Nature 559, 217–222.

Li, R., Bitoun, E., Altemose, N., Davies, R.W., Davies, B., and Myers, S.R. (2019a). A high-resolution map of non-crossover events reveals impacts of genetic diversity on mammalian meiotic recombination. Nat. Commun. 10, 1–15.

Li, W., Yang, L., He, Q., Hu, C., Zhu, L., Ma, X., Ma, X., Bao, S., Li, L., Chen, Y., et al. (2019b). A Homeostatic Arid1a-Dependent Permissive Chromatin State Licenses Hepatocyte Responsiveness to Liver-Injury-Associated YAP Signaling. Cell Stem Cell 25, 54–68.e5.

Long, H., Zhang, L., Lv, M., Wen, Z., Zhang, W., Chen, X., Zhang, P., Li, T., Chang, L., Jin, C., et al. (2020). H2A.Z facilitates licensing and activation of early replication origins. Nature 577, 576–581.

Maezawa, S., Yukawa, M., Alavattam, K.G., Barski, A., and Namekawa, S.H. (2018). Dynamic reorganization of open chromatin underlies diverse transcriptomes during spermatogenesis. Nucleic Acids Res. 46, 593–608.

Marchal, C., Sima, J., and Gilbert, D.M. (2019). Control of DNA replication timing in the 3D genome. Nat. Rev. Mol. Cell Biol. 20, 721–737.

Miotto, B., Ji, Z., and Struhl, K. (2016). Selectivity of ORC binding sites and the relation to replication timing, fragile sites, and deletions in cancers. Proc. Natl. Acad. Sci. U. S. A. 113, E4810–E4819.

Monesi, V. (1962). Autoradiographic study of DNA synthesis and the cell cycle in spermatogonia and spermatocytes of mouse testis using tritiated thymidine. J. Cell Biol. 14, 1–18.

Murakami, H., and Keeney, S. (2014). Temporospatial coordination of meiotic DNA replication and recombination via DDK recruitment to replisomes. Cell 158, 861–873.

Murakami, H., and Nurse, P. (2001). Regulation of premeiotic S phase and recombination-related double-strand DNA breaks during meiosis in fission yeast. Nat. Genet. 28, 290–293.

Murakami, H., Lam, I., Huang, P.-C., Song, J., van Overbeek, M., and Keeney, S. (2020). Multilayered mechanisms ensure that short chromosomes recombine in meiosis. Nature 582, 124–128.

Myers, S., Bottolo, L., Freeman, C., McVean, G., and Donnelly, P. (2005). A fine-scale map of recombination rates and hotspots across the human genome. Science 310, 321–324.

Myers, S., Bowden, R., Tumian, A., Bontrop, R.E., Freeman, C., MacFie, T.S., McVean, G., and Donnelly, P. (2010). Drive against hotspot motifs in primates implicates the PRDM9 gene in meiotic recombination. Science 327, 876–879.

Neale, M.J., Pan, J., and Keeney, S. (2005). Endonucleolytic processing of covalent protein-linked DNA double-strand breaks. Nature 436, 1053–1057.

Parvanov, E.D., Petkov, P.M., and Paigen, K. (2010). Prdm9 controls activation of mammalian recombination hotspots. Science 327, 835.

Patel, L., Kang, R., Rosenberg, S.C., Qiu, Y., Raviram, R., Chee, S., Hu, R., Ren, B., Cole, F., and Corbett, K.D. (2019). Dynamic reorganization of the genome shapes the recombination landscape in meiotic prophase. Nat. Struct. Mol. Biol. 1.

Petryk, N., Kahli, M., d’Aubenton-Carafa, Y., Jaszczyszyn, Y., Shen, Y., Silvain, M., Thermes, C., Chen, C.-L., and Hyrien, O. (2016). Replication landscape of the human genome. Nat. Commun. 7, 10208.

Petryk, N., Dalby, M., Wenger, A., Stromme, C.B., Strandsby, A., Andersson, R., and Groth, A. (2018). MCM2 promotes symmetric inheritance of modified histones during DNA replication. Science 361, 1389–1392.

Picard, F., Cadoret, J.-C., Audit, B., Arneodo, A., Alberti, A., Battail, C., Duret, L., and Prioleau, M.-N. (2014). The spatiotemporal program of DNA replication is associated with specific combinations of chromatin marks in human cells. PLoS Genet. 10, e1004282.

Powers, N.R., Parvanov, E.D., Baker, C.L., Walker, M., Petkov, P.M., and Paigen, K. (2016). The Meiotic Recombination Activator PRDM9 Trimethylates Both H3K36 and H3K4 at Recombination Hotspots In Vivo. PLoS Genet. 12, e1006146.

Pratto, F., Brick, K., Khil, P., Smagulova, F., Petukhova, G.V., and Camerini-Otero, R.D. (2014). Recombination initiation maps of individual human genomes. Science 346, 1256442.

Quinlan, A.R., and Hall, I.M. (2010). BEDTools: a flexible suite of utilities for comparing genomic features. Bioinformatics 26, 841–842.

Ramírez, F., Dündar, F., Diehl, S., Grüning, B.A., and Manke, T. (2014). deepTools: a flexible platform for exploring deep-sequencing data. Nucleic Acids Res. 42, W187–W191.

R Core Team (2014). R: A language and environment for statistical computing (R Foundation for Statistical Computing, Vienna, Austria.).

Sandhu, R., Monge Neria, F., Monge Neria, J., Chen, X., Hollingsworth, N.M., and Börner, G.V. (2020). DNA Helicase Mph1FANCM Ensures Meiotic Recombination between Parental Chromosomes by Dissociating Precocious Displacement Loops. Dev. Cell 53, 458–472.e5.

Sequeira-Mendes, J., Vergara, Z., Peiró, R., Morata, J., Aragüez, I., Costas, C., Mendez-Giraldez, R., Casacuberta, J.M., Bastolla, U., and Gutierrez, C. (2019). Differences in firing efficiency, chromatin, and transcription underlie the developmental plasticity of the Arabidopsis DNA replication origins. Genome Res. 29, 784–797.

Smagulova, F., Gregoretti, I.V., Brick, K., Khil, P., Camerini-Otero, R.D., and Petukhova, G.V. (2011). Genome-wide analysis reveals novel molecular features of mouse recombination hotspots. Nature 472, 375–378.

Smagulova, F., Brick, K., Pu, Y., Camerini-Otero, R.D., and Petukhova, G.V. (2016). The evolutionary turnover of recombination hot spots contributes to speciation in mice. Genes Dev. 30, 266–280.

Smith, D.I., Zhu, Y., McAvoy, S., and Kuhn, R. (2006). Common fragile sites, extremely large genes, neural development and cancer. Cancer Lett. 232, 48–57.

Smith, O.K., Kim, R., Fu, H., Martin, M.M., Lin, C.M., Utani, K., Zhang, Y., Marks, A.B., Lalande, M., Chamberlain, S., et al. (2016). Distinct epigenetic features of differentiation-regulated replication origins. Epigenetics Chromatin 9, 18.

Stinchcomb, D.T., Thomas, M., Kelly, J., Selker, E., and Davis, R.W. (1980). Eukaryotic DNA segments capable of autonomous replication in yeast. Proc. Natl. Acad. Sci. U. S. A. 77, 4559–4563.

Takahashi, S., Miura, H., Shibata, T., Nagao, K., Okumura, K., Ogata, M., Obuse, C., Takebayashi, S.-I., and Hiratani, I. (2019). Genome-wide stability of the DNA replication program in single mammalian cells. Nat. Genet. 51, 529–540.

Valton, A.-L., Hassan-Zadeh, V., Lema, I., Boggetto, N., Alberti, P., Saintomé, C., Riou, J.-F., and Prioleau, M.-N. (2014). G4 motifs affect origin positioning and efficiency in two vertebrate replicators. EMBO J. 33, 732–746.

Walker, M., Billings, T., Baker, C.L., Powers, N., Tian, H., Saxl, R.L., Choi, K., Hibbs, M.A., Carter, G.W., Handel, M.A., et al. (2015). Affinity-seq detects genome-wide PRDM9 binding sites and reveals the impact of prior chromatin modifications on mammalian recombination hotspot usage. Epigenetics Chromatin 8, 31.

Wu, P.-Y.J., and Nurse, P. (2014). Replication origin selection regulates the distribution of meiotic recombination. Mol. Cell 53, 655–662.

Wu, H., Caffo, B., Jaffee, H.A., Irizarry, R.A., and Feinberg, A.P. (2010). Redefining CpG islands using hidden Markov models. Biostatistics 11, 499–514.

Yevshin, I., Sharipov, R., Valeev, T., Kel, A., and Kolpakov, F. (2017). GTRD: a database of transcription factor binding sites identified by ChIP-seq experiments. Nucleic Acids Res. 45, D61–D67.

Yin, Y., Jiang, Y., Lam, K.-W.G., Berletch, J.B., Disteche, C.M., Noble, W.S., Steemers, F.J., Camerini-Otero, R.D., Adey, A.C., and Shendure, J. (2019). High-Throughput Single-Cell Sequencing with Linear Amplification. Mol. Cell.

Zhang, Y., Liu, T., Meyer, C.A., Eeckhoute, J., Johnson, D.S., Bernstein, B.E., Nusbaum, C., Myers, R.M., Brown, M., Li, W., et al. (2008). Model-based analysis of ChIP-Seq (MACS). Genome Biol. 9, R137.

